# A neural basis of choking under pressure

**DOI:** 10.1101/2023.04.16.537007

**Authors:** Adam L. Smoulder, Patrick J. Marino, Emily R. Oby, Sam E. Snyder, Hiroo Miyata, Nick P. Pavlovsky, William E. Bishop, Byron M. Yu, Steven M. Chase, Aaron P. Batista

## Abstract

Incentives tend to drive improvements in performance. But when incentives get too high, we can “choke under pressure” and underperform when it matters most. What neural processes might lead to choking under pressure? We studied Rhesus monkeys performing a challenging reaching task in which they underperform when an unusually large “jackpot” reward is at stake. We observed a collapse in neural information about upcoming movements for jackpot rewards: in the motor cortex, neural planning signals became less distinguishable for different reach directions when a jackpot reward was made available. We conclude that neural signals of reward and motor planning interact in the motor cortex in a manner that can explain why we choke under pressure.

**One-Sentence Summary:** In response to exceptionally large reward cues, animals can “choke under pressure”, and this corresponds to a collapse in the neural information about upcoming movements.

## Main Text

Failing to perform to one’s highest standard when the potential payoff is particularly great is known as “choking under pressure” (*1*). While failures in professional athletics often provide the most memorable examples of this phenomenon, people also choke under pressure in a wide variety of other settings, including test-taking, video games, puzzle-solving, and more (*2–7*).

Neuroimaging studies have implicated the involvement of reward and motor structures in choking under pressure (*8–10*), but the neural mechanisms whereby the possibility of increased rewards can lead to performance failure remain unclear.

We recently reported that animals also choke under pressure (*11*). Rhesus monkeys performed a challenging task in which they had to perform a goal-directed reach that was both fast and accurate (**Fig. 1A**). We cued the animals as to the magnitude of the liquid reward they would receive for a successful reach. Performance in the task was influenced by reward size: success was more likely for Medium and Large potential rewards than for Small rewards. This presumably reflects the motivation to perform this challenging task. However, success rates fall when “Jackpot” (rare and exceptionally large) rewards are proffered, leading to the “inverted-U” relationship between performance and reward that characterizes choking under pressure (**Fig. 1B**). Here we leverage the fact that monkeys choke under pressure to explore the phenomenon’s neural basis at the resolution of the activity of individual neurons and the sub-second timescale at which neural activity controls behavior.

**Fig. 1.**
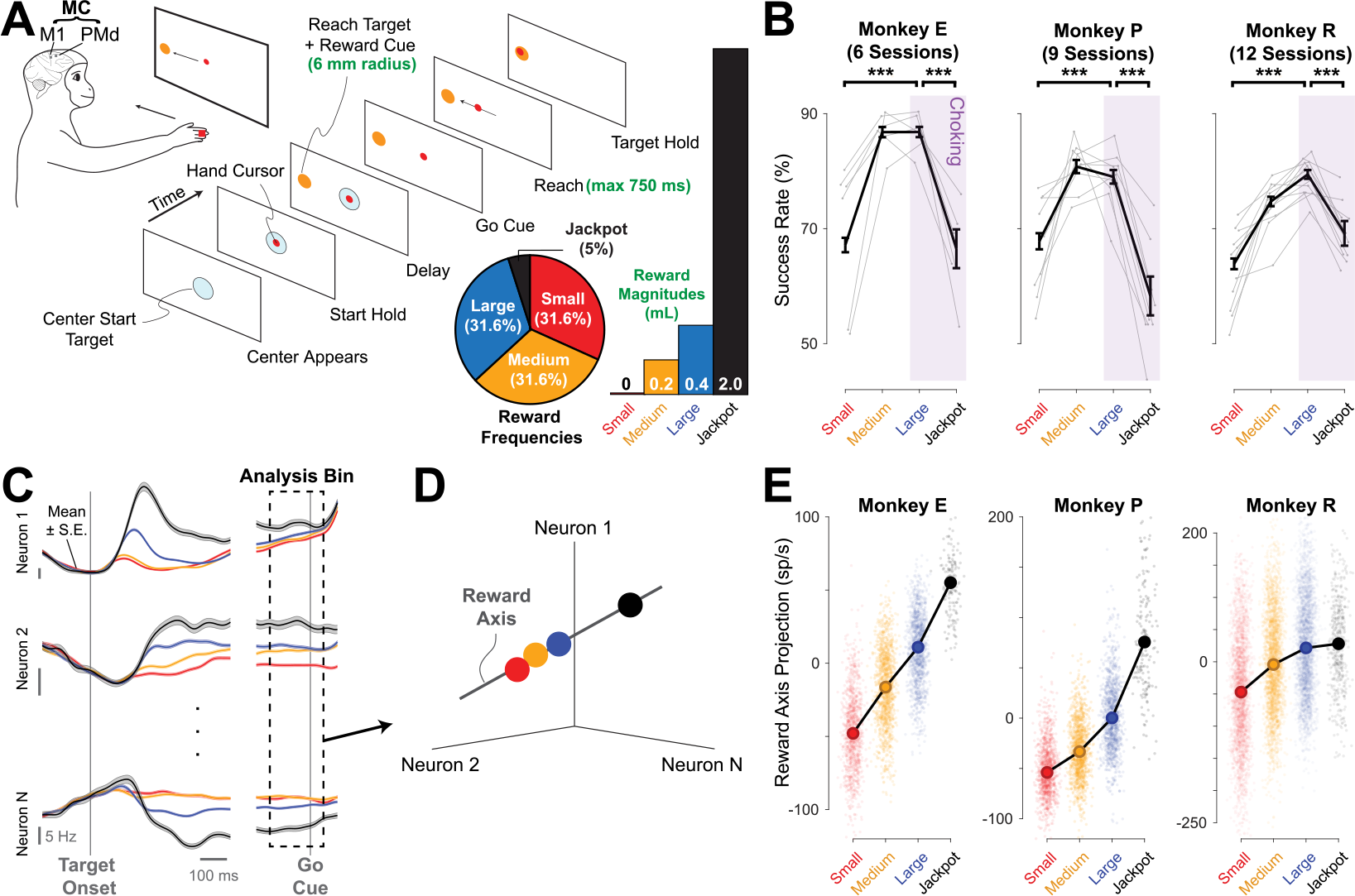
Monkeys choke under pressure, although reward tuning in motor cortex is monotonic. **(A)** Monkeys were trained to prepare then reach briskly to a small target. The color (Monkeys E, P) or shape (Monkey R) instructed the reward size. Parameters bolded in green were titrated for each animal to make the task challenging and motivating (**Table S1** shows task parameters for each animal). A separate choice task indicated that animals understood reward cues (**Fig. S1**). Simultaneously, we recorded from primary motor (M1) and/or dorsal premotor cortex (PMd) using 96-channel microelectrode “Utah” arrays (Blackrock Microsystems, Inc., gray squares schematize general array locations, see **Table S1** for array location details). **(B)** Success rate improved from Small to Large rewards (binomial proportion test, ***p < 0.001), indicating that performance in this difficult task is influenced by motivation. All three animals choked under pressure, indicated by the significant decrease in task success rates from Large to Jackpot rewards. Error bars represent S.E. of overall mean success rate shown in black. Individual sessions are shown in gray. **(C)** Individual neurons exhibited monotonic tuning to reward size. Activity traces from three example neurons from Monkey E are shown, averaged within each reward condition (± S.E.). We highlight the time window used for the ensuing neural analyses: during reach preparation at the end of the delay period, a time when the animal had information regarding both the target location and potential reward size to be received for a successful trial. **(D)** Simultaneous neural firing rates can be visualized in a neural state space in which the firing rate of each neuron comprises one dimension (axis) within the space. Three neurons were used here for illustration; in actuality, hundreds of neurons recorded over 6-12 days were used (see Methods for the stitching procedure used to combine data across sessions). **(E)** The dimension that captures the majority of reward-related variance follows monotonic trends with cued reward, even though behavior exhibits a non-monotonic relationship with reward. Translucent dots show single trial values, while the large, filled dots show the mean of the reward condition. A horizontal jitter is introduced within each group to improve visibility. We considered whether these reward-monotonic effects might reflect muscular stiffening. However, neither arm nor shoulder electromyography showed much activity during the reach planning period, and neither reliably predicted reward axis projections (**Fig. S2**).

We report a novel neural explanation of choking under pressure: a deficit in motor planning. Motor planning benefits the execution of rapid, voluntary movements (*12, 13*), like the reaches the animals performed in this task. To study how motor planning relates to choking under pressure, we recorded the spiking activity of neurons in the motor cortex (MC, the primary motor cortex and the dorsal aspect of the premotor cortex) and examined how the cued reward modulated neural population activity during movement planning. MC sends the predominant cortical projection to the spinal cord for the control of arm movements and encodes information about planned movements (*14–17*). If choking under pressure involves a failure in motor planning, we might expect there to be aspects of MC activity that exhibit an inverted-U relationship with reward size, like behavior does.

We first asked how the magnitude of the cued reward affected the firing rate of individual neurons in MC. Neural signals of anticipated reward have been reported throughout the cerebral cortex, with neurons in many brain areas exhibiting changes in firing rates when more valuable rewards are cued (*18–25*), including neurons in MC (*26–28*). However, previous studies have not presented monkeys with rare and exceptionally large potential rewards that induce performance decrements, and thus the nature of the cortical response to such Jackpot rewards is unknown. Given the “inverted-U” profile that characterizes how behavioral performance is impacted by reward size, it is reasonable to ask whether the encoding of reward by individual neurons also follows the inverted-U profile.

The reward tuning in MC was predominantly monotonic. The majority of MC neurons exhibited tuning to cued reward (n = 300/459 neurons, 65.4%; single neuron metrics are provided in full in **Table S2**), where most exhibited either monotonically increasing (179/459, 39.0%) or decreasing (95/459, 20.7%) changes in firing rate through the entire range of cued reward size (**Fig. 1C**, see Methods). We observed little “inverted-U” (18/459, 3.9%) or “U-shaped” (8/459, 1.7%) reward tuning in firing rates. Thus, we conclude that although individual MC neurons are sensitive to Jackpot reward cues, the basis for the inverted-U profile evident in behavior is not to be found in the reward-driven changes in the average activity of individual MC neurons.

Next, we considered whether neural signatures of choking under pressure might be present at the population level. We analyzed patterns of covariance in the activity of simultaneously recorded neurons. By treating the activity of each individual neural unit as an axis in a high dimensional space, we can identify specific dimensions (i.e., linear combinations of neurons’ firing rates) that capture reward-related variance (**Fig. 1D**). Because there were so few Jackpots given per session, for analyses we combined neural activity across days using a “stitching” algorithm (*29*) (see Methods). We then used principal components analysis (PCA) on trial-averaged activity to identify the linear projection maximizing the amount of reward-related variance captured. We found that a single dimension, which we call the “reward axis,” captured the majority of the reward-related variance (Monkey E: 92.6%, P: 89.8%, R: 84.7%). Consistent with the single-neuron responses, projections along the reward axis were monotonic with reward size (**Fig. 1E**). In sum, we find that the encoding of reward information in MC is primarily monotonic, which on its own is not able to explain the performance drop observed for Jackpot rewards.

Since we did not see evidence for choking under pressure in the neural signal of reward considered on its own, we next wondered whether an interaction might exist between reward signals and the neural activity associated with motor planning. Individual neurons in MC are tuned for different directions of upcoming reaches, such that at the neural population level, distinct neural activity patterns correspond to the motor plans for different reach directions (*14, 17*). We hypothesized that reward information may interact with the directional reach planning signals in MC, and that this interaction might lead to choking under pressure.

To look for such an interaction, we began by identifying the neural subspace that contained reach direction signals. We found a projection of the trial-averaged neural population activity using PCA that provided the maximal separation of average neural activity corresponding to different motor plans. The top 2 principal components accounted for the overwhelming majority of the variance due to target direction (Monkey E: 92.7%, P: 99.7%, R: 90.8%). This plane turned out to be nearly orthogonal to the reward axis (**Fig. S3**). Even with this near orthogonality, however, we observed an interaction between reward cue and target direction. Comparing neural activity for Small, Medium, and Large cues, the mean response for the different upcoming movement directions grew farther apart from one another with increased reward **(Fig. 2A**) (*30*). This can reflect greater information about the upcoming reach with larger rewards, as the average neural activity patterns for different movements are more distinct from one another. Surprisingly, for Jackpot rewards, the neural states for different target directions collapsed towards each other (*31*).

**Fig. 2.**
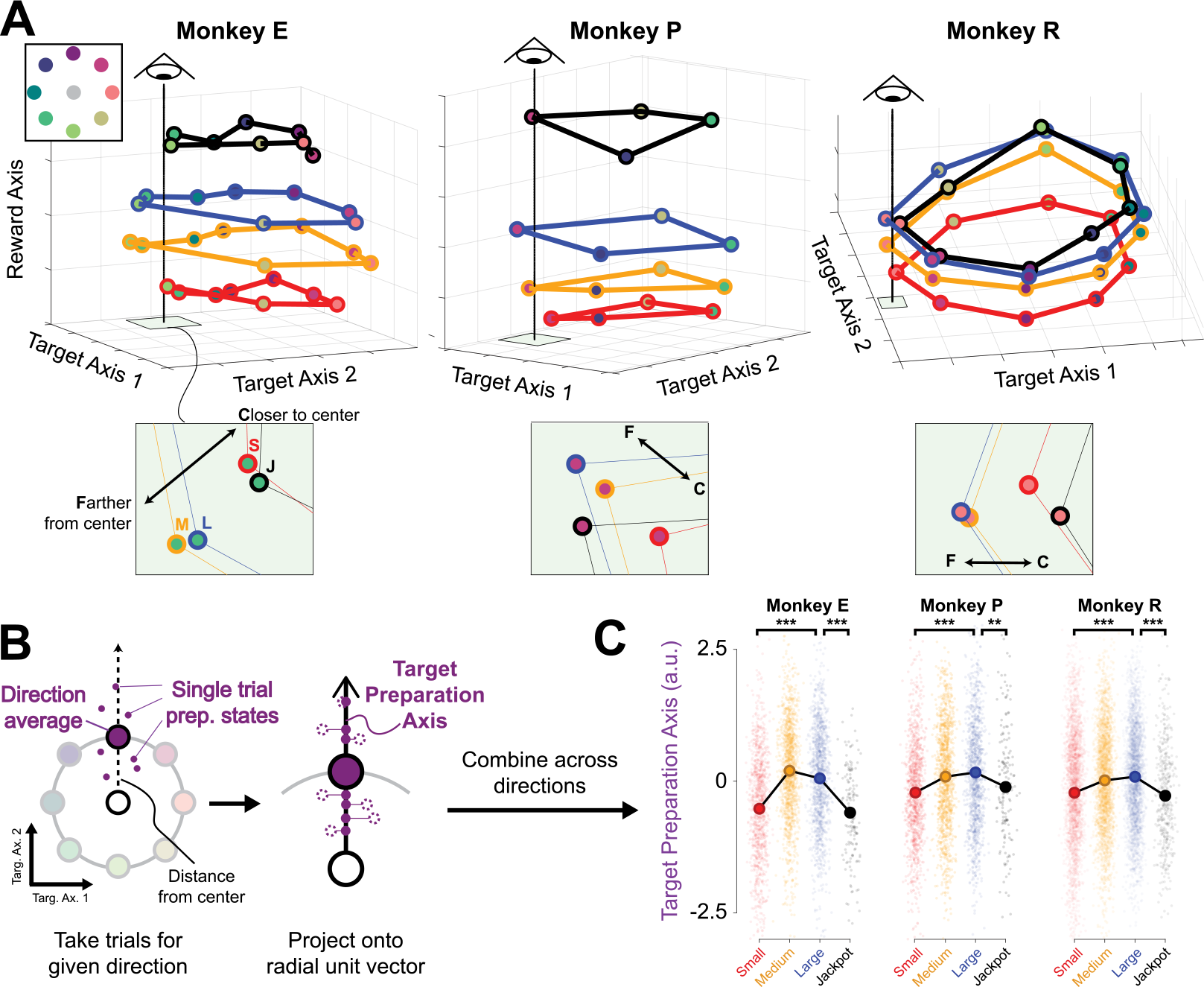
Jackpot rewards induce a collapse in neural information. This is evident in the interaction between the neural population response for movement direction and reward. **(A)** Neural population activity corresponding to motor planning for different reach directions is pushed apart with increasing cued reward from Small through Large. However, for Jackpot rewards, the activity for different reach directions collapses back towards each other, diminishing their discriminability. We projected neural activity grouped by trial conditions defined by reward and direction and then averaged into a 3D space reflecting reward information (Reward Axis) and target information (Target Axis 1 and 2). The units (population neural activity, spikes/s) on the three axes are the same. To aid visualization, adjacent reach directions (dot color) are connected by a ring for each reward (line color). Insets show a zoomed-in view of the target axes’ plane for a single target to highlight the inverted-U interaction of reward and direction on neural activity. **(B)** To quantify single trial separability of preparatory states, we found a “target preparation axis” for each reward and target direction (see Methods). **(C)** When neural activity for individual trials is projected onto these target preparation axes, it exhibits an inverted-U as a function of cued reward that parallels the behavior. Dots represent single trials, and large filled circles show the mean within each reward condition. **p < 0.01, ***p < 0.001, Welch’s t-test.

To quantify this expansion-then-collapse of neural states with reward, we examined neural activity on individual trials. Within each reward condition, we first identified the average response for each target direction (large purple dot in **Fig. 2B**) and calculated the average across the targets (large white dot in **Fig. 2B**). We then constructed unit vectors that pointed to each target’s average from the average response across targets. We call these vectors the “target preparation axes”. We projected the neural activity for each trial onto the corresponding target preparation axis. Like success rates, the average projection along the target preparation axis follows an inverted-U as a function of reward (**Fig. 2C**), congruent with the visualizations from Figure 2A. We refer to this decrease from Large to Jackpot rewards as a collapse in neural information (*32*). That is, *target* information becomes less discriminable as neural population activity moves along the *reward* axis from Large to Jackpot states. In this manner, the neural population activity resembles the animal’s behavior, in that both show an inverted-U dependence on reward size.

How might a collapse in neural information be connected to a decrease in behavioral performance? We hypothesized that when the neural state was further out along the target preparation axis, this might correspond to better reach preparation. Thus, a collapse in neural information, indicated by small projections onto the target preparation axis, would correspond to poorer preparation of the reach (**Fig. 3A**). We compared the magnitude of the projection of neural activity onto the target preparation axis to the animals’ performance. We separated the trials into successes and failures and then we categorized failed trials by their specific failure mode. The animals could fail by executing a reach that either overshoots or undershoots the target (**Fig. 3B**; see Methods). The decrease in success rate between Large and Jackpot reward trials was dominated by undershoot failures (**Fig. 3C**) (*11, 33*). We conclude from this analysis that the collapse in neural information when Jackpot rewards are proffered coincides with undershooting the target.

**Fig. 3.**
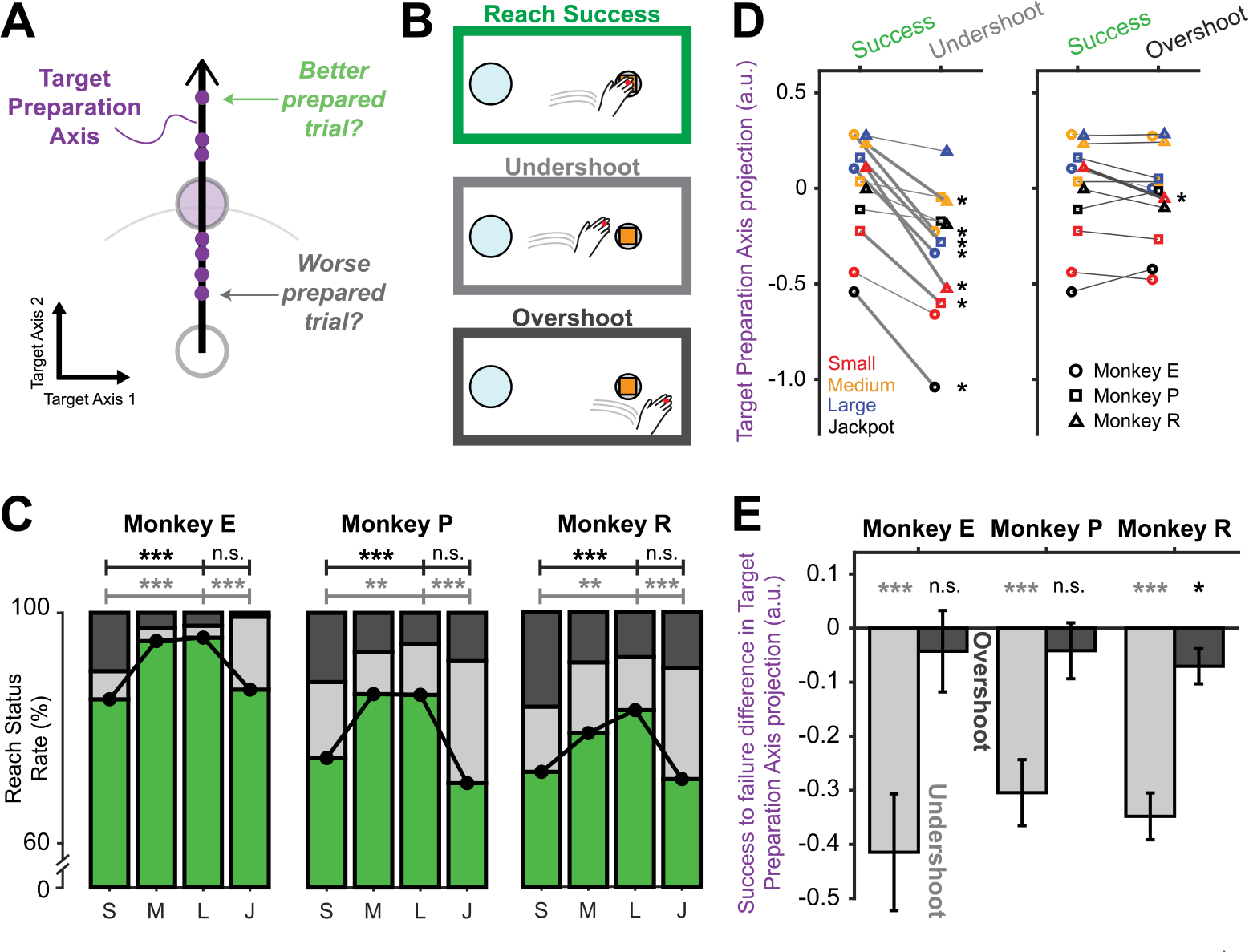
The quality of reach preparation is reflected in the state of neural activity. **(A)** Hypothesized relationship between single trial target preparation axis projections (small purple dots) and reach preparation. **(B)** Possible reach outcomes. Monkeys can either succeed at the reach (top), undershoot the target (middle), or overshoot the target (bottom). **(C)** Success rates (green) and failure rates broken down by failure type (light gray: undershoots; dark gray: overshoots). Compared to Large rewards, Jackpots had more undershoots (binomial proportion test; for all panels, *p < 0.05, **p < 0.01, ***p < 0.001, n.s. = not significant) but not more overshoots, whereas Small rewards evoked both more undershoots and overshoots than did Large rewards. **(D)** Undershoots showed a consistent decrease in average target preparation axis projections across animals (shape) and rewards (color) compared to successes, whereas overshoot trials showed little difference in their projection along the target preparation axis. Thick lines and accompanying stars indicate a significant difference within that given reward condition (Welch’s t-test). **(E)** To summarize the relationship between target preparation axis projections and failure modes, we pooled across rewards after z-scoring based on successful trials within each (mean ± S.E.). Trials that result in undershoots (left, light gray) show a significant decrease in projections along the target preparation axis (Welch’s t-test), while overshoot failures (right, dark gray) show a much smaller effect.

As a more stringent test of this relationship, we examined it on individual trials. We projected neural activity onto the target preparation axis (see Methods) and labeled it according to whether the trial was a success, a failure due to an undershoot, or a failure due to overshooting the target. Within every reward condition, neural preparatory activity prior to an undershoot failure had a smaller projection along the target preparation axis than preparatory activity prior to a success (**Fig. 3D**, *left*). In contrast, there was little difference between neural activity on overshoot failures and successes (**Fig. 3D**, *right*). This means that when the projection of neural activity onto the target preparation axis was smaller for a given trial, the animal was more likely to fail by undershooting the target. Quantifying this across all trials for each animal revealed that undershoot trials had significantly smaller target preparation axis projections than successes (**Fig. 3E**). This effect also holds when using other algorithms to define the target preparation axis (**Fig. S8**). This observation links the collapse in neural information triggered by a Jackpot reward to the decline in behavioral performance for Jackpots. We suggest that choking under pressure is due to an adverse interaction of reward information with movement preparation signals, and that this interference is visible in motor cortex.

Our analyses so far support the view that one neural basis of choking under pressure is due to a poor positioning of neural activity relative to an optimal region that lies further outward along the target preparation axis. We also considered another explanation for choking under pressure: that it is due to variability in neural activity. Note that reward could in principle affect both the position of neural population activity in the neural state space and also its variability, so this effect could occur alongside the changes in average activity reported above.

Variability in neural activity across trials in motor cortical planning activity is known to be a major source of variability in behavior (*34–36*). Neural variability can depend on context; as an example, songbirds are known to modulate their amount of neural variability during song production depending on whether they are practicing their song alone or performing for courtship (*37*). Hence it could be that choking under pressure results from an increase in neural variability induced by the Jackpot reward cue. To look for an explanation of choking under pressure stemming from reward-induced effects on variability, we calculated trial-to-trial variability at the population level. We found inconsistent relationships between neural variability and reward across our three subjects, and no evidence for a U-shaped relationship between reward and neural variability (**Fig. S9**). Hence our data do not support an explanation for choking under pressure in terms of neural variability.

In summary, we can describe a potential neural basis for choking under pressure: Reward information interacts with the formation of motor command signals. This interaction can be seen in planning-related neural activity in the motor cortex. Reward information can help boost neural information (evident in the transition from Small to Large rewards). But when a Jackpot is proffered, neural activity does not attain the optimal preparation state for a well-executed movement. The specific way in which these states are suboptimal is that they are less differentiated according to the upcoming reach target. That is, a “collapse in neural information” occurs when a Jackpot is proffered, and this corresponds to a decrease in performance. These poor planning states are correlated with the propensity to fail by undershooting the target. In broader scope, our findings are a striking example of context altering movement preparatory activity and the ensuing input-output transformation implemented by motor cortex (*38–44*).

Choking under pressure is a robustly observed phenomenon across many forms of cognitive, sensorimotor, and perceptual tasks with multiple potential psychological explanations (*2, 6, 8, 9, 45–47*). Studies of humans implicate many brain areas in choking under pressure, including the basal ganglia, prefrontal cortex, and motor cortex (*8–10*). This suggests that the neural bases of choking are widespread in the brain, perhaps reflecting the action of neuromodulators. Our study shows a candidate neural mechanism for choking under pressure in motor cortex where an interaction between information about the reward and behaviorally relevant neural signals corresponds to under-performance when the stakes are unusually high. The neural basis for choking under pressure we report here might be a specific example of a widespread phenomenon: it may be that choking under pressure is the result of adverse interactions between motivational signals and diverse neural functions, including cognition and perception, leading to a collapse in the neural information supporting various types of behavior.

## Acknowledgments

We thank Simon Borgognon for feedback on the manuscript, Hongwei Mao for assistance in experimental setup, and Alan Degenhart for contributing to task design.

## Funding

Achievement Rewards for College Scientists, Pittsburgh Chapter Award (ALS)

Achievement Rewards for College Scientists, Abraham–Martin–Ragni Award (NPP)

Bradford and Diane Smith Graduate Fellowship in Engineering (ALS)

National Science Foundation graduate research fellowship DGE1745016 (ALS)

National Science Foundation graduate research fellowship DGE2139321 (PJM)

Howard Hughes Medical Institute (WEB)

National Science Foundation grant BCS 1533672 (SMC, BMY, APB)

National Science Foundation grant DRL2124066 (BMY, SMC) / DRL2123911 (APB)

National Institutes of Health grant R01HD071686 (APB, SMC, BMY)

National Institutes of Health CRCNS grant R01NS105318 (BMY, APB)

National Institutes of Health grant R01NS129584 (APB, SMC, BMY)

National Institutes of Health grant R01NS129098 (APB, SMC)

Simons Foundation grant 543065 (BMY)

## Author contributions

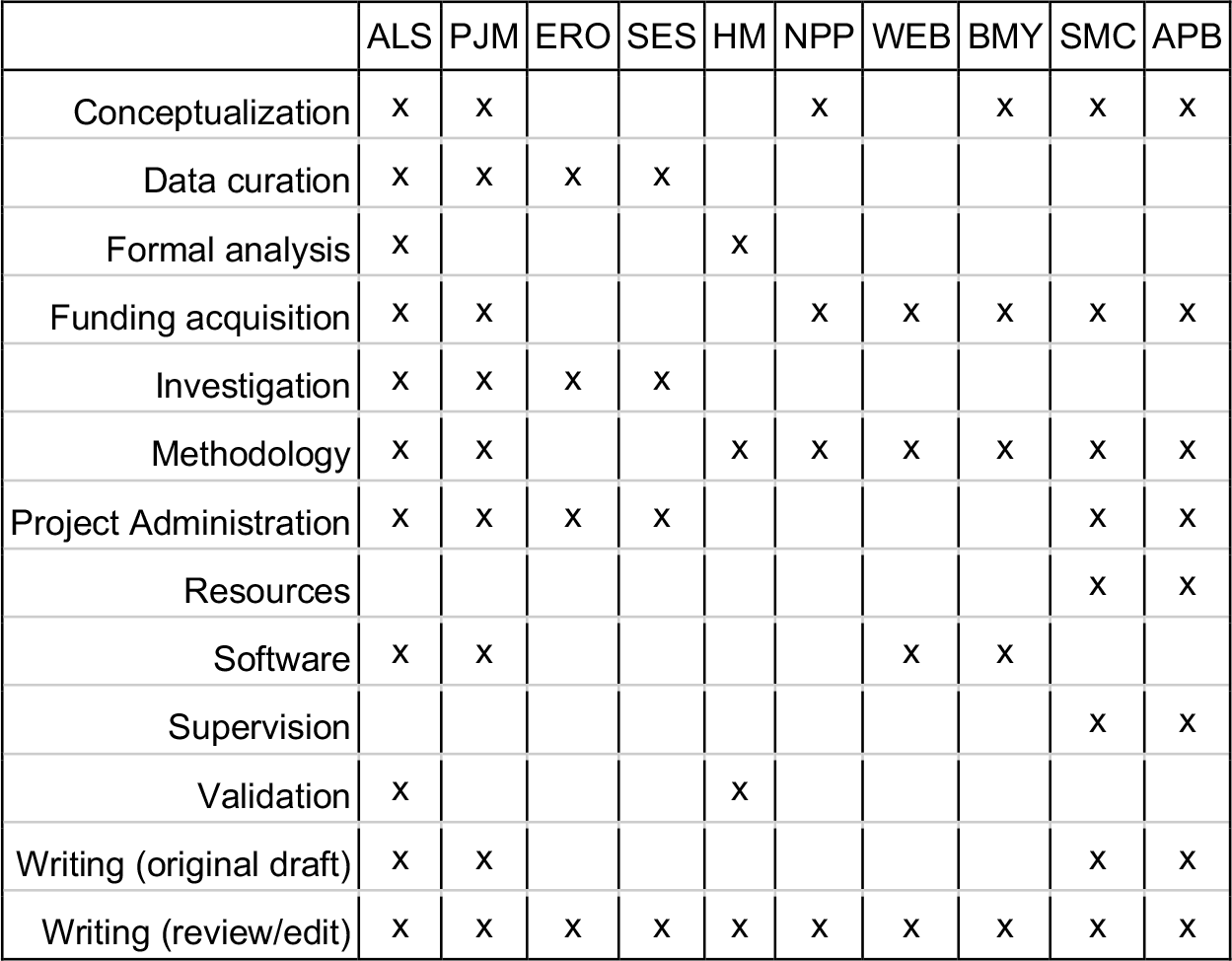

## Data and materials availability

Data will be made available at the time of publication.

## Supplementary Materials

Materials and Methods

Figs. S1 to S10

Tables S1 to S2

References (48-58)

Movie S1

## References and Notes

1. Although there appears to be no technical term better-suited than this expression from everyday parlance, similar terms are found around the world, such as “cracking”, “weakening”, “folding”, “wavering”, or “collapsing” under pressure. This paradoxical decrease in performance when payoffs or stakes increase seems ubiquitous.

2. R. F. Baumeister, Choking under pressure: Self-consciousness and paradoxical effects of incentives on skillful performance. Journal of Personality and Social Psychology, 11 (1984).

3. C. E. Kimble, J. S. Rezabek, Playing games before an audience: Social facilitation or choking. Soc. Behav. Pers. 20, 115–120 (1992).

4. S. L. Beilock, T. H. Carr, When high-powered people fail. Psychological Science. 16, 5 (2005).

5. D. F. Gucciardi, J. A. Dimmock, Choking under pressure in sensorimotor skills: Conscious processing or depleted attentional resources? Psychology of Sport and Exercise. 9, 45–59 (2008).

6. B. P. Lewis, D. E. Linder, Thinking about choking? Attentional processes and paradoxical performance. Personality and Social Psychology Bulletin. 23, 937–944 (1997).

7. D. Ariely, U. Gneezy, G. Loewenstein, N. Mazar, Large stakes and big mistakes. Review of Economic Studies. 76, 451–469 (2009).

8. T. G. Lee, S. T. Grafton, Out of control: Diminished prefrontal activity coincides with impaired motor performance due to choking under pressure. NeuroImage. 105, 145–155 (2015).

9. V. S. Chib, B. De Martino, S. Shimojo, J. P. O’Doherty, Neural mechanisms underlying paradoxical performance for monetary incentives are driven by loss aversion. Neuron. 74, 582–594 (2012).

10. V. S. Chib, S. Shimojo, J. P. O’Doherty, The effects of incentive framing on performance decrements for large monetary outcomes: behavioral and neural mechanisms. Journal of Neuroscience. 34, 14833–14844 (2014).

11. A. L. Smoulder, N. P. Pavlovsky, P. J. Marino, A. D. Degenhart, N. T. McClain, A. P. Batista, S. M. Chase, Monkeys exhibit a paradoxical decrease in performance in high-stakes scenarios. Proc. Natl. Acad. Sci. U.S.A. 118, e2109643118 (2021).

12. A. M. Haith, J. Pakpoor, J. W. Krakauer, Independence of movement preparation and movement initiation. Journal of Neuroscience. 36, 3007–3015 (2016).

13. K. C. Ames, S. I. Ryu, K. V. Shenoy, Simultaneous motor preparation and execution in a last-moment reach correction task. Nat Commun. 10, 2718 (2019).

14. J. Tanji, E. V. Evarts, Anticipatory activity of motor cortex neurons in relation to direction of an intended movement. Journal of Neurophysiology. 39, 1062–1068 (1976).

15. D. J. Crammond, J. F. Kalaska, Prior information in motor and premotor cortex: activity during the delay period and effect on pre-movement activity. Journal of Neurophysiology. 84, 986–1005 (2000).

16. M. M. Churchland, G. Santhanam, K. V. Shenoy, Preparatory activity in premotor and motor cortex reflects the speed of the upcoming reach. Journal of Neurophysiology. 96, 3130–3146 (2006).

17. N. Even-Chen, B. Sheffer, S. Vyas, S. I. Ryu, K. V. Shenoy, Structure and variability of delay activity in premotor cortex. PLoS Comput Biol. 15, e1006808 (2019).

18. M. Watanabe, Reward expectancy in primate prefrontal neurons. Nature. 382 (1996).

19. L. Tremblay, W. Schultz, Relative reward preference in primate orbitofrontal cortex. Nature. 398, 704–708 (1999).

20. M. L. Platt, P. W. Glimcher, Neural correlates of decision variables in parietal cortex. Nature. 400, 233–238 (1999).

21. L. P. Sugrue, Corrado, G. S., Newsome, W. T., Matching behavior and the representation of value in the parietal cortex. Science. 304, 1782–1787 (2004).

22. S. Musallam, B. D. Corneil, B. Greger, H. Scherberger, R. A. Andersen, Cognitive control signals for neural prosthetics. Science. 305, 258–262 (2004).

23. B. Y. Hayden, J. M. Pearson, M. L. Platt, Fictive reward signals in the anterior cingulate cortex. Science. 324, 948–950 (2009).

24. S. W. Kennerley, T. E. J. Behrens, J. D. Wallis, Double dissociation of value computations in orbitofrontal and anterior cingulate neurons. Nat Neurosci. 14, 1581–1589 (2011).

25. L. Stănişor, C. van der Togt, C. M. A. Pennartz, P. R. Roelfsema, A unified selection signal for attention and reward in primary visual cortex. Proc. Natl. Acad. Sci. U.S.A. 110, 9136–9141 (2013).

26. M. R. Roesch, C. R. Olson, Impact of expected reward on neuronal activity in prefrontal cortex, frontal and supplementary eye fields and premotor cortex. Journal of Neurophysiology. 90, 1766–1789 (2003).

27. M. R. Roesch, C. R. Olson, Neuronal activity related to reward value and motivation in primate frontal cortex. Science. 304, 307–310 (2004).

28. B. T. Marsh, V. S. A. Tarigoppula, C. Chen, J. T. Francis, Toward an autonomous brain machine interface: Integrating sensorimotor reward modulation and reinforcement learning. Journal of Neuroscience. 35, 7374– 7387 (2015).

29. A. D. Degenhart, W. E. Bishop, E. R. Oby, E. C. Tyler-Kabara, S. M. Chase, A. P. Batista, B. M. Yu, Stabilization of a brain–computer interface via the alignment of low-dimensional spaces of neural activity. Nat Biomed Eng. 4, 672–685 (2020).

30. Movie S1 shows rotations of the space in Fig. 2; Fig. S4 shows similar results using algorithms other than PCA to identify target encoding dimensions.

31. The same effect of expansion and then collapse in neural encoding with reward can also be observed in single-unit tuning curves (Fig. S5).

32. The technical definition of information is a reduction in uncertainty, which incorporates both changes in condition averages (“signal”) and variability (“noise”). While Figure 2 focuses on changes in signal, noise will be analyzed later in the paper. To back up the claim that these changes in signal reflect a collapse in neural information, we used an offline decoding approach. This analysis found that decode accuracy also exhibited the inverted-U (Fig. S6).

33. Other types of errors were also more present for Jackpot trials, but those other failure modes were not consistent across animals. Undershoot failures can be caused by slow reaction, slow peak reach speed, and a shorter reach plan. The animals show a mixture of these factors contributing to an increased amount of undershoot failures occurring for both Small and Jackpot trials compared with Large (Fig. S7).

34. M. M. Churchland, A. Afshar, K. V. Shenoy, A central source of movement variability. Neuron. 52, 1085–1096 (2006).

35. K. S. Chaisanguanthum, H. H. Shen, P. N. Sabes, Motor variability arises from a slow random walk in neural state. Journal of Neuroscience. 34, 12071–12080 (2014).

36. P. O. Boucher, T. Wang, L. Carceroni, G. Kane, K. V. Shenoy, C. Chandrasekaran, “Neural population dynamics in dorsal premotor cortex underlying a reach decision” (preprint, Neuroscience, 2022),, doi:10.1101/2022.06.30.497070.

37. S. C. Woolley, R. Rajan, M. Joshua, A. J. Doupe, Emergence of context-dependent variability across a basal ganglia network. Neuron. 82, 208–223 (2014).

38. M.-C. Hepp-Reymond, M. Kirkpatrick-Tanner, L. Gabernet, H.-X. Qi, B. Weber, Context-dependent force coding in motor and premotor cortical areas. Experimental Brain Research. 128, 123–133 (1999).

39. R. G. Rasmussen, A. Schwartz, S. M. Chase, Dynamic range adaptation in primary motor cortical populations. eLife. 6, e21409 (2017).

40. S. Naufel, J. I. Glaser, K. P. Kording, E. J. Perreault, L. E. Miller, A muscle-activity-dependent gain between motor cortex and EMG. Journal of Neurophysiology. 121, 61–73 (2019).

41. J. A. Hennig, E. R. Oby, M. D. Golub, L. A. Bahureksa, P. T. Sadtler, K. M. Quick, S. I. Ryu, E. C. Tyler-Kabara, A. P. Batista, S. M. Chase, B. M. Yu, Learning is shaped by abrupt changes in neural engagement. Nat Neurosci. 24, 727–736 (2021).

42. S. Vyas, M. D. Golub, D. Sussillo, K. V. Shenoy, Computation through neural population dynamics. Annu. Rev. Neurosci. 43, 249–275 (2020).

43. X. Sun, D. J. O’Shea, M. D. Golub, E. M. Trautmann, S. Vyas, S. I. Ryu, K. V. Shenoy, Cortical preparatory activity indexes learned motor memories. Nature. 602, 274–279 (2022).

44. C. F. Fisac, S. M. Chase, “Sensory constraints on volitional modulation of the motor cortex” (preprint, Neuroscience, 2023),, doi:10.1101/2023.01.22.525098.

45. R. M. Yerkes, J. D. Dodson, The relation of strength of stimulus to rapidity of habit-formation. J. Comp. Neurol. Psychol. 18, 459–482 (1908).

46. J. A. Easterbrook, The effect of emotion on cue utilization and the organization of behavior. Psychological Review. 66, 183–201 (1959).

47. J. Wine, Test anxiety and direction of attention. Psychological Bulletin. 76, 92–104 (1971).

48. M. D. Golub, B. M. Yu, S. M. Chase, Internal models for interpreting neural population activity during sensorimotor control. eLife. 4, e10015 (2015).

49. W. E. Bishop, B. M. Yu, “Deterministic symmetric positive semidefinite matrix completion” in Advances in Neural Information Processing Systems (2014).

50. C. Pandarinath, D. J. O’Shea, J. Collins, R. Jozefowicz, S. D. Stavisky, J. C. Kao, E. M. Trautmann, M. T. Kaufman, S. I. Ryu, L. R. Hochberg, J. M. Henderson, K. V. Shenoy, L. F. Abbott, D. Sussillo, Inferring single-trial neural population dynamics using sequential auto-encoders. Nat Methods. 15, 805–815 (2018).

51. J. Jude, M. G. Perich, L. E. Miller, M. H. Hennig, Robust alignment of cross-session recordings of neural population activity by behaviour via unsupervised domain adaptation (2022), (available at http://arxiv.org/abs/2202.06159).

52. W. E. Bishop, thesis, Carnegie Mellon University (2015).

53. M. Nonnenmacher, S. C. Turaga, J. H. Macke, “Extracting low-dimensional dynamics from multiple large-scale neural population recordings by learning to predict correlations” in Advances in Neural Information Processing Systems (2017).

54. G. W. Fraser, A. B. Schwartz, Recording from the same neurons chronically in motor cortex. Journal of Neurophysiology. 107, 1970–1978 (2012).

55. J. A. Gallego, M. G. Perich, R. H. Chowdhury, S. A. Solla, L. E. Miller, Long-term stability of cortical population dynamics underlying consistent behavior. Nat Neurosci. 23, 260–270 (2020).

56. B. M. Yu, J. P. Cunningham, G. Santhanam, S. I. Ryu, K. V. Shenoy, M. Sahani, Gaussian-process factor analysis for low-dimensional single-trial analysis of neural population activity. Journal of Neurophysiology. 102, 614–635 (2009).

57. A. Georgopoulos, J. Kalaska, R. Caminiti, J. Massey, On the relations between the direction of two-dimensional arm movements and cell discharge in primate motor cortex. J. Neurosci. 2, 1527–1537 (1982).

58. S. Bates, T. Hastie, R. Tibshirani, Cross-validation: what does it estimate and how well does it do it? (2022), (available at http://arxiv.org/abs/2104.00673).

