## Supplemental Materials for "A neural basis of choking under pressure"

---------------------------------------------------------------------------------------------

**The PDF file includes:**

Materials and Methods
Figs. S1 to S10
Tables S1 to S2

**Other Supplementary Materials for this manuscript include the following:**

Movie S1

### Materials and Methods

Experiments and behavioral recordings

All experimental and animal procedures were approved by the University of Pittsburgh Institutional Animal Care and Use Committee in accordance with the guidelines of the US Department of Agriculture, the International Association for the Assessment and Accreditation of Laboratory Animal Care, and the National Institutes of Health. All analyses were performed using MATLAB R2021a.

*Task setup and kinematic recordings*

Three adult male rhesus macaques, Monkeys E (9.0 kg, 10 years old), P (9.5 kg, 6 years old), and R (19.0 kg, 10 years old) were trained on delayed reaching tasks. We used standard water regulation procedures to maintain motivation and the valuation of reward. During experiments, each monkey sat in a primate chair facing a mirror ~8 cm in front of his eyes that reflected a computer monitor displaying task events. The monkeys performed all tasks by making hand movements (right arm for Monkeys E and R, left arm for Monkey P) in an open space in front of them. While the working arm and hand were unrestrained, the hand was not visible to the animal, as it moved in the space behind and below the mirror. 3D hand position was tracked with an infrared LED marker attached either to the monkey's index finger (Monkeys E and P, 120 Hz sampling rate, nominal resolution < 1 mm, PhaseSpace, Inc) or the back of the hand (Monkey R, 60 Hz sampling rate, nominal resolution < 1 mm, Optotrak 3020, Northern Digital Instruments). The monkey's hand movements corresponded to a cursor position displayed on the monitor. The software environment was calibrated such that 1 cm of hand displacement in the coronal plane corresponded to 1 cm of cursor movement. Any trials where tracking of the hand failed at any point in time were removed (fewer than 1% of trials).

For post-hoc analysis, we smoothed the hand position signals with a zero-phase low pass Butterworth filter (total 8th order, cutoff frequency 15 Hz). We then also calculated velocity by taking the first difference of the position data, dividing by the time difference between samples, and assigning each velocity sample a timestamp that was the midpoint of the timestamps for the two position samples used. We then used spline interpolation for both the position and velocity signals to upsample to 1000 Hz at identical timepoints. For Monkey R, we also simultaneously recorded surface electromyography (EMG) from shoulder and arm muscles (see Electromyography section below).

*Main task*

The main task performed by the animals in this study was a challenging delayed center out reaching task (**Fig. 1A**) which has been described in detail in (*11*) (referred to as the "speed + accuracy task"). Specific task parameters for each animal are listed in **Table S1**. All trials began with a circular target appearing at the center of the display. The animal had to move its hand to position the cursor in the target to initiate a trial. After a short period of time (ranging from 200-600 ms, different for each animal), a reach target appeared at one of several possible locations (Monkeys E and R: 8 possible target locations, Monkey P: 4) and remained visible through the rest of the trial. The appearance of the reach target began a delay period during which the animal could see their hand position cursor, the reach target, and the center target, but was not allowed to let the cursor exit the center target. The duration of this delay period was drawn randomly from a uniform distribution each trial (ranging from 200-950 ms, different distributions for each animal). After this duration, the center target disappeared from the screen, which cued the animal to move the cursor to the reach target. The animal had only a brief amount of time after the center target disappeared to acquire the reach target (range across animals of 667-825 ms). If the animal moved the cursor into the reach target in time, he had to hold the cursor within the target for 400 ms, after which he received a liquid reward. The allowed reach duration and target size were titrated during training for each animal to make the task challenging, and then those values were maintained throughout experiments.

The magnitude of the liquid reward that would be dispensed upon successful completion of the trial was indicated by the color of the target cue (Monkeys E, P) or by an image within the target (Monkey R). Four reward sizes were used for each animal and were drawn randomly every trial: Small, Medium, Large, and Jackpot (magnitudes and cues differed by animal; see **Table S1)**. Jackpot rewards were presented on only 5% of trials (which resulted in a range of 1-12 Jackpots proffered per reach direction per day), while the other rewards were presented with equal frequency for the remaining trials. For purposes of visualization in this work, we use the color scheme indicated in **Fig. 1A**: Small is red, Medium is orange, Large is blue, and Jackpots are black (see **Table S1** for actual cue colors and shapes).

For Monkeys P and R, a small proportion of trials (5%) were "catch" trials, where the trial abruptly ended at the time of go cue and the animal was rewarded without making a reach. These were used to discourage "false start" reaches before the go cue. Catch trials were randomly interspersed throughout the session and could be of any cued reward and direction. The reward provided on catch trials corresponded to the cued reward.

*Two target choice task*

To test whether the animals understood the reward cues, we instructed them to perform a two-target choice (**Fig. S1A)**. This task proceeded in the same way as the main task with two differences: (1) two reach targets were available for the animal to select, diametrically opposed, and (2) the animal had a longer amount of time to reach to their selected target. Target direction was selected randomly from the working animal's target set used in the main task. Reward values were typically selected randomly each trial with equal probability, with the exception that Jackpot rewards were available for selection only on 5% of trials and the Jackpot reward was never presented at both available targets. Upon trial success, the animal would receive the reward associated with the target he selected.

We then evaluated the animals' selections in each combination of reward presentations to confirm the animals selected the greater reward when presented. Collectively, animals selected the larger reward on 98.9% of trials (**Fig. S1B**). Trials where both targets had the same reward cue were excluded from this analysis.

In both the main and two-target task, the animal failed the trial if he moved the hand position cursor outside of the center target during the delay period, did not acquire the reach target in time, or exited the reach target before 400 ms had elapsed after acquiring it. Upon failure, the cursor and reach target turned purple and the screen froze for a duration exceeding the maximum length remaining for a successful trial (i.e., if the animal failed during the delay epoch, the freeze time was the remaining time for the longest delay period, plus the maximum reach time, plus the 400 ms target hold). After this, a purple reach target appeared at a location other than that of the failed target. The animal had to reach to the purple target, for which no reward was given, before the next trial could begin. These "punishment" reaches were included to discourage the animal from aborting Small reward trials, as it makes task failure objectively worse than attempting the trial, even for the least rewarding (e.g., 0 mL) cases. After either a punishment reach was completed (for failed trials), or a reward was dispersed (for successful trials), an intertrial interval elapsed before the next trial began (Monkeys E and P: 500 ms, Monkey R: 600 ms). For Monkey R, intertrial intervals after successful Jackpot trials were extended to 1200 ms to give additional time for the animal to consume the larger reward.

*Behavioral training and recording sessions*

All animals followed the same general training procedures prior to performing the task for the recordings used in this study. Animals were first trained to proficiently perform a delayed reaching task (> 90% success rate). Then, reward cues for Small, Medium, and Large were introduced into this task. We ran sessions of the two-target choice task after the main task each day to assess understanding of the relative reward cue values. After at least one week of performing the same task with these reward cues, we began making the task more challenging. We titrated the amount of time the animal had to acquire the reach target and the reach target size to reduce average success rate to about 70%. Making the task challenging enabled us to better measure if cued reward either improved or hurt success rate. If the difference in success rate between Small and Large rewards was small (i.e., less than 10%), we expanded the reward range by shrinking the Small reward magnitude and increasing the Large reward. We expanded the Small-Large reward range in this manner for two animals; Monkey R before introduction of Jackpot rewards, and Monkeys E after Jackpots had been introduced (see (*11*)).

Once the animals indicated understanding of the relative value of the reward cues in the choice task and parameters were adjusted to the appropriate task difficulty, we introduced the Jackpot reward cue. For each animal, we ran the main task for one session, then the next day split the session between the main task and the two-target choice task to assess understanding of the Jackpot cue. Monkey E indicated full Jackpot cue understanding, after which we performed experiments. After several sessions of this, we then expanded Monkey E's Small-Large reward range and ran the choice task each day after the main task. These sessions with the expanded Small-Large reward range are what is reported in this study and are shown in **Fig. S1B** (see (*11*) for information on the initial main task sessions). Monkey P indicated full Jackpot cue understanding during this session, after which we halted Jackpot behavioral experiments until after array implantation. After recovery, we ran a session of choice task including Jackpots to ensure Monkey P still understood cue values (the data of which are shown in **Fig. S1B**), after which we collected data for the main task. Monkey R did not immediately select the Jackpot cue when presented, potentially because of a more challenging cue to learn and distinguish (the cue was a specific image, as opposed to a color). Additionally, his performance improved in the first 2-3 sessions after Jackpot reward introduction. We adjusted the parameters of the task to account for these behaviors (we shortened the reach period maximum time but slightly increased reach target diameter). After this, we repeated the procedure of one session with all four reward cues and one session of split main and choice task, during which the animal indicated understanding of all reward cues. We began neurophysiological recordings the following day. The data from this session of the choice task and additionally run choice task sessions after a few days of the main task are reported in **Fig. S1B**.

Behavioral analysis

*Success rate and failure mode analysis*

We analyzed success rates for the main task as a function of reward cue (**Fig. 1B**). Average success rates were pooled across all sessions (black lines), and also calculated for individual sessions (gray lines). For error bars, we performed a bootstrap analysis in which we randomly sampled (with replacement) the trials within each reward size and calculated the success rate 10,000 times. Error bars are the standard deviation of these 10,000 bootstrap samples.

We analyzed the way the animals failed trials in the same manner as previous work (*11*). For this study, we only considered trials where the go cue occurred (i.e., no failure during the delay period). Once the go cue occurred, two types of reach failures could occur: overshoots and undershoots (**Fig. 3B**). First, the animal could reach past the target, miss it, and run out of time prior to being able to make a correction, an "overshoot." We also considered trials where the cursor blew through the target as overshoot failures. Second, the animal could "undershoot" by running out of time before acquiring the target. Undershoots could occur either due to a reach landing short of the target and the animal having insufficient time for corrective movements, or due to reacting and/or reaching slowly and running out of time mid-reach. Although these two types of undershoots are potentially distinguishable based on kinematics, we reasoned that both could correspond to a failure to plan well, and thus we combined them for analyses. If the cursor stopped successfully in the target, it was considered a reach epoch success (a success for the purposes of **Fig. 3C**); however, the animal could still fail if the cursor drifted out of the reach target during the target hold period (a failure for the purposes of overall success rate in **Fig. 1B**). The specific parameters used to decide which label each failed trial was given are described in (*11*). We note that the failure mode trends in this study for comparing Small and Jackpot reward trials to Large reward trials were consistent with the results from (*11*).

*Single trial behavioral metrics*

We report four kinematic metrics as a function of reward in **Fig. S7**. To find peak speed for each trial, we first calculated the cursor speed as the square root of the sum of the squared horizontal and vertical velocity signals. We calculated the peak speed of each trial as the max cursor speed occurring after the go cue. We calculated the reaction time as the time from the go cue that it took for the cursor to reach 20% of the animal's overall average peak speed. In this way, reaction time was calculated at the crossing of a static speed threshold for each animal.

We calculated the homing time and ballistic endpoint predictions in the same way as previous work (*11*). Homing time was calculated starting at the time the cursor was within the last ⅓ of the distance to the target and ending when the cursor was within 1 mm of the target edge. Homing time trends were not sensitive to this exact start or end point. Ballistic endpoint predictions were calculated by first identifying the locations of the cursor 50 ms before reaction time and at the time of peak speed. The displacement between these points was calculated and doubled. This is mathematically equivalent to mirroring the velocity profile in time about peak speed and integrating, producing a prediction of where the reach would have landed with a symmetric, ballistic velocity profile. We report this quantity specifically projected along the vector connecting the center target to the reach target.

### Electromyography

For one animal (Monkey R), we recorded surface electromyography (EMG) on various muscles of the working arm and its shoulder.

*Surface EMG recordings*

We used pairs of disposable adhesive electrodes for bipolar recordings (3M, model 2560), placed with a dab of conductive gel (Spectra 360) above muscles on shaved regions of the working arm (lateral biceps, triceps) and shoulder (anterior deltoid, posterior deltoid, trapezius). Anecdotally, shoulder muscle signals were more salient than those from arm muscles. We also placed a pair of electrodes across the chest to record electrocardiogram (ECG) signals. A ground electrode was placed on the skin above the lower, outer stomach. Signals were sent through a differential amplifier (RA16LI-D, Tucker Davis Technologies (TDT), Inc.), then a pre-amplifier (PZ2 Pre-Amp, TDT), then to a signal processor that digitized signals (nominal 48KHz sampling frequency, RZ2 BioAmp Processor, TDT). Signals were then fed through a 2nd order 10 Hz biquad high pass filter, downsampled to 10KHz and stored. Two minutes of data were collected before the start of each experiment to identify baseline EMG and ECG.

*Surface EMG preprocessing*

For analysis, EMG and ECG data were zero-phase filtered using a comb of notch filters to remove line noise and a bandpass Butterworth filter to attenuate movement artifacts and high-frequency noise. The comb filter was 2$^{nd}$ order, with cutoff frequencies of 60 Hz and its harmonics up to 480 ± 1 Hz. The bandpass filter had cutoff frequencies of 20 and 500 Hz with stopband attenuation of -60 dB, implemented using MATLAB's "bandpass" function. ECG artifacts were visible in the EMG signals during recording, particularly during the two minute baseline period. To remove these, we fit a 50 ms transfer function from the ECG data to each of the EMG recordings using the two-minute baseline period, then applied this transfer function to the entire EMG recordings. This produced a predicted ECG artifact signal for each EMG channel, which was subtracted out, after which signals were rectified and downsampled to 1000 Hz. To combine signals across days, we z-scored data each day using the mean and standard deviation of a directional "tuning curve" calculated using the average EMG activation in a 200 ms window surrounding peak cursor speed (see Behavioral analysis), with an additional point added to the tuning curve to include EMG in the 200 ms period before target onset while Monkey R held the cursor at the center target.

*Surface EMG analysis*

We performed two analyses on Monkey R's surface EMG signals. First, we assessed the overall quality of the EMG signals by visualizing activity for each reach direction (**Fig. S2A**). For all successful trials, we smoothed rectified EMG signals with a 50 ms boxcar filter and extracted three windows aligned to target onset, go cue, and target acquisition. We then averaged across all trials within each reach direction and plotted activity as a function of reach direction for each muscle.

Second, we assessed EMG during the delay period time bin coinciding with the neural analyses in the main text ([-150 50] ms around the time of go cue) to determine if muscle activation could explain changes in neural activity with increased cued rewards. For any trial that had a go cue (i.e., no delay failures or catch trials) we calculated the average EMG signal in this time bin to get one value per muscle per trial, then grouped trials by their reward and direction labels. We then performed a two-way ANOVA to evaluate statistical significance of reward and directional tuning in the EMG signals (**Fig. S2B**).

Neural recordings and preprocessing

We recorded neural activity from the primary motor cortex (M1) and dorsal aspect of the premotor cortex (PMd) using multielectrode "Utah" arrays (Blackrock Microsystems). Monkey E had one 96-electrode array straddling the shoulder regions of M1 (~64 channels) and PMd (~32 channels), implanted approximately 1.5 years prior to these experiments. Monkey P had two 64 electrode arrays, one in the arm area of M1 and one in the arm area of PMd, implanted two weeks prior to these experiments. Due to equipment constraints, only 96 electrodes could be recorded simultaneously at the time of the experiments, so only half of the M1 electrodes were used. Monkey R had two 96 electrode arrays in the anterior and posterior aspects of M1, in the shoulder region, implanted approximately 4.5 years prior to these experiments. Recordings on most electrodes from both regions showed modulation during the late delay period analyzed here, and thus in this study we make no distinction between recordings from M1 versus PMd, referring to the combined site of recordings as motor cortex (MC). We include all recordings for all analyses.

Before each experiment began, we set voltage thresholds on each electrode at a multiplier of the root-mean-square voltage (Monkey E: -3.5x, P: -3x, R: -3x). We stored 1 ms waveform snippets (sampling rate 30 KHz; 30 samples per snippet) surrounding each threshold crossing during the experiment. Individual units were manually sorted on each electrode using offline spike sorting (Plexon). Units were identified visually using a combination of features, including waveform principal components and peak-to-trough amplitude. We included both well-isolated units and multi-unit waveforms for this study, yielding each day 145±4 units for Monkey E (mean ± std across sessions), 183±86 units for Monkey P, and 273±25 units for Monkey R.

Neural data analysis

In this work, we focused on steady-state neural activity during reach preparation, a 200 ms period starting 150 ms before each trial's go cue and ending 50 ms after. As 50 ms is less than the time it takes for visual information to reach MC (*48*), we consider all of this activity to fall within the time the animal was prepared to move but had yet to begin reacting to the go cue. We calculated the averaged firing rate within this bin. For the visualizations of **Fig. 1C**, we made peri-stimulus time histograms for the delay period using spike times convolved with a Gaussian kernel (25 ms standard deviation).

Monkeys P and R experienced a subset of trials with very short delay periods (< 400 ms). To assess if neural activity sufficiently reached a steady state for these short-delay trials, we calculated the average distance of the neural state from a steady state endpoint (~450 ms after target onset) as a function of time after target onset for non-short delay trials (*17*). We found that Monkey E's trajectories reached 90% of the average distance by 220 ms, P by 197 ms, and R by 317 ms. We excluded trials with delays shorter than this for all analyses. We note that this procedure only removes trials for Monkey R (shortest delay 200 ms), as Monkey E and P's shortest delay period is longer than this. Using a static threshold of trials with delay length greater than 400 ms did not meaningfully affect results.

*Single unit analysis*

For single unit analyses (**Table S2, Fig. S5**), we only included units that were present for at least 10 presentations of Jackpot rewards for each reach direction (leaving 42 units for Monkey E, 113 units for Monkey P, and 304 units for Monkey R). This typically required the unit to be present across at least 2-3 sessions of data collection (see *Combining neural data across* *sessions*). To analyze tuning as a function of reach direction and reward, we used a 2-way analysis of variance (ANOVA) on the binned firing rates for each identified unit. If the ANOVA indicated significant effects of reward on firing rate, we used Tukey's test with multiple comparisons correction (the "multcompare" function in MATLAB) to assess significant differences between reward conditions, from which we could evaluate the shape of the reward tuning curve. For this analysis only, we combined Medium and Large rewards due to their general similarity in effects on firing rate and to simplify the operation of evaluating reward tuning curve shape. Based on the pattern of the differences between the Small, Medium/Large, and Jackpot reward conditions, we identified 9 possible tuning curve shapes, as described in **Table S2**. We categorized each unit into one of these 9 shapes and tallied their results into **Table S2**.

*Combining neural activity across sessions*

Jackpot rewards occurred rarely (5% of trials), meaning that single sessions have a low quantity of Jackpot trials for a given reach direction. Combining across days is necessary to gain the statistical power sufficient to analyze the Jackpot trials. However, due to electrode recording instabilities, not all units are conserved session-to-session, and few neural units are present across all recording sessions (21 units for Monkey E, 3 units for Monkey P, and 6 units for Monkey R; see below for details about how this was assessed). Statistical algorithms can be used to "stitch" neural recordings across sessions (*49–53*). In this work, we combined neural activity across days using a modified version of the stabilization algorithm from Degenhart, Bishop et al. 2020 (*29*), described below. It requires identification of common units across recording sessions but does not require consistent task or trial structure across sessions. Unless stated otherwise, all further methods and analyses were performed using the latent factors identified by this neural stitching method.

First, to identify units that were present in recordings across different sessions, we used the method described in (*54*). In brief, we calculated four quantities for every sorted unit using the first 100 trials of each session: (1) average waveform shape, (2) average firing rate, (3) spiking autocorrelation, and (4) spiking cross-correlations with all other sorted units from that session. We then compared these four quantities between all possible pairs of units on different sessions and calculated similarity scores for each quantity for each possible pair of units between the sessions as described in (*54*). For instance, if session 1 had $n_1$ recorded units and session 2 had $n_2$, we made a total of $n_1 * n_2$ comparisons between those sessions. We assumed that units recorded from different electrodes were not the same across days due to the physical distance between electrodes; these different channel unit comparisons were used to compose a "different" distribution for each of the four similarity scores. The similarity scores for pairs of units on the same channel across days were compared to this distribution. If the likelihood of the similarity scores being from the "different" distribution was below a false-positive threshold (0.01 for Monkeys E and R, and 0.05 for Monkey P), the units were considered the same. Using this procedure, we found a high number of units tracked over different pairs of recording sessions, primarily across consecutive sessions. For example, the minimum number of units present across two consecutive sessions was 71 for Monkey E, 28 for Monkey P, and 67 for Monkey R. Units could be present across non-consecutive sessions (i.e., present sessions 1 and 3 but not session 2). However, this was rare and nearly exclusively occurred for Monkey P, whose array was implanted shortly preceding the study and, as such, exhibited greater amounts of instability in recordings.

Having identified and tracked units across sessions, we organized the neural activity into a $n \times T$ matrix of binned spike counts, where $n$ is the number of unique units recorded (i.e., the union of units across all sessions), and $T$ is the total number of trials across all sessions. If unit $i$ was present during trial $t$, that element of the matrix was filled in with the spike count. Otherwise, that element was left empty.

To identify a common latent space across sessions (the $T$ trials), we used factor analysis (FA). We chose FA because it is the most basic dimensionality reduction method that seeks to preserve variance shared across units. FA relates neural activity for trial $t$, $\boldsymbol{x_t} \in \mathbb{R}^n$, to latent factors, $\boldsymbol{z_t} \in \mathbb{R}^m$ ($m < n$) according to:

$$\boldsymbol{x_t} | \boldsymbol{z_t} \sim N(\Lambda \boldsymbol{z_t} + \boldsymbol{\mu}, \Psi) \qquad (1)$$

$$\boldsymbol{z_t} \sim N(\mathbf{0}, I) \qquad (2)$$

where $\Lambda \in \mathbb{R}^{n \times m}$ is the loading matrix that defines the relationship between latent factors and spike counts, $\boldsymbol{\mu} \in \mathbb{R}^n$ is a vector of mean spike counts for each unit, and $\Psi \in \mathbb{R}^{n \times n}$ is a diagonal matrix that describes the variability independent to each unit. The latent factors $\boldsymbol{z_t}$ capture the variance that is shared across units.

To fit the FA model parameters $\Lambda$, $\Psi$, and $\boldsymbol{\mu}$, we used the expectation maximization (EM) algorithm. Notably, there are empty entries in the spike count vectors, $\boldsymbol{x_t}$, as described above. We treat this as a missing data problem, where units that are not present for a given session are treated as missing observations. The EM algorithm can handle missing data seamlessly by maximizing the probability of only the data entries that were observed. By fitting a single FA model across the $T$ trials, we effectively "stitch" the neural activity across experimental sessions to identify a single common latent space. This method is described in (*52*), with theory that can be used to derive how many units need to be in common across recordings for successful stitching in (*49*). Other methods are also available for neural stitching, for example those that leverage the time course of activity (*51*, *53*) or trial labels such as reach direction (*55*). Note that the stitching method we use here does not require trial labels; instead, it leverages the identification of stable units across (a subset of) sessions.

To determine the number of factors $m$, we used 5-fold cross validation to fit FA models ranging from 1 to 40 factors. We selected the number of factors $m^*$ that maximized the cross-validated data likelihood (Monkey E: 16 factors, P: 5, R: 20). We then fit a single FA model for each animal across all observed activity $\boldsymbol{x_t}$ using $m^*$ factors. This procedure was performed separately for each animal, yielding a single "stitched" FA model per animal. To facilitate visualizations and analyses involving the latent factors, we orthonormalized the columns of the loading matrix $\Lambda$ and ordered the factors (i.e., the elements of $\boldsymbol{z_t}$) by the amount of shared variance explained (*56*). We refer to the latent factors simply as "neural activity" for the remainder of the methods and in the main text from Fig. 1E onward.

To validate the stitching procedure, we assessed the stability of neural direction and reward representations across days (**Fig. S10**); more rigorous validation of the algorithm itself is available in (*52*). Whenever possible, we reproduced results found in the stitched latent space with analyses on single units (**Table S2, Fig. S5**).

*Reward axis calculation*

To find a population-level signature of reward encoding in the motor cortex, we identified a signal that captured reward-related variance across the population. We first calculated the trial-averaged neural activity within each direction and reward condition. We then marginalized across reach directions, yielding a number-of-rewards (4) by number-of-factors matrix of average neural activity for each reward. We performed principal components analysis (PCA) on this matrix to identify the dimensions explaining the most reward-related variance. A single PC explained the overwhelming majority of the reward-related variance for each animal (E: 92.6%, P: 89.8%, R: 84.7%); because of this, we dubbed this PC the "reward axis". Since PCA output is only unique to a sign-flip, we selected the sign of the reward axis such that Large reward trials had greater average projections than Small reward trials.

We projected all trials along this single dimension for each animal (**Fig. 1E**). We note that there was no part of the PCA objective that encouraged monotonic trends for Jackpot rewards; that is, if the main reward-related variance in the data exhibited an inverted-U, it would have come out of this procedure. We found near-identical reward axis results when we used other algorithms to identify it, such as linear discriminant analysis (LDA) with reward labels or linear regression of neural activity to categorical reward size (i.e., S = 1, M = 2, L = 3, J = 4, data not shown).

*Target axes calculation*

We wanted to study how reward interacted with the representation of target direction during movement planning. To determine the dimensions in neural space with the most target-related variance, we once again calculated the trial-averaged activity within each direction and reward condition, then marginalized across rewards to produce a number-of-directions by number-of-factors matrix for each animal. We then performed PCA on this to find the linear combinations of factors that capture the greatest variance about target response. We found that the top two PCs explained the overwhelming majority of this variance (E: 92.7%, P: 99.7%, R: 90.8%). (Note that this large amount of variance explained by two dimensions for a static time bin of neural data is not particularly surprising: the number of points used is equal to the number of directions, and accordingly the number of dimensions needed to fully account for these points' variance is the number of targets minus one. Further, the target positions were embedded in a 2D space in the coronal plane, meaning only two dimensions are needed to fully define the target locations.) We called the axes of this 2D space "target axes". For ease of visualization after projecting to this 2D plane, we aligned all animal's spaces such that the average neural activity for the up-rightward reach target was pointed along the first quadrant diagonal unit vector (meaning that target axis 1 is positive for rightward targets, while target axis 2 is positive for upward targets).

*Projecting data into a 3D space made of the target and reward axes*

The target axes are orthogonal in the latent space by definition when using PCA. However, as the reward axis and the target axes were found in separate steps, there were no constraints to the alignment of the reward axis with respect to the target axes. We assessed how aligned the reward and target axes were for each animal and found the reward axis to be near-orthogonal to the target axes' plane (**Fig. S3**). This permits orthogonalization of the reward axis to the target plane with minimal distortion of reward axis projection values.

Given this near-orthogonality, for purposes of visualization only (i.e., **Fig. 2, Fig. S4, Fig. S5**), we orthogonalize the reward axis with respect to the 2D target plane using QR decomposition.

We projected all trials into this 3D space, then calculated the average activity within each direction and reward condition (**Fig. 2A**). We reproduced the results shown in **Fig. 2A** using other algorithms to identify the target axes, including LDA with target direction labels and linear regression of neural activity to spatial target location (**Fig. S4A**).

*Target preparation axes*

We saw qualitatively that the separation of average activity for different upcoming reach directions (i.e., the radius of the target ring) was modulated as a function of cued reward in **Fig. 2A**. To quantify this effect, we performed an analysis on single trials calculating how close or far the neural preparatory state was from the origin of the target axes' plane. To do this, we calculated "target preparation axes" within each reward and direction condition as follows (**Fig. 2B**): First, we calculated the average projections for each reward and direction condition in the target axes' plane. Then, for a given reward condition, we calculated the average across directions. Intuitively, this is the center of the ring of targets for the given reward shown in **Fig. 2A**. Then, for a given reward-direction condition, we calculated the vector connecting this ring center for the given reward to the average activity for the current reward-direction. We then projected all trials' activity for that condition along the vector. Higher values indicate greater distance from the ring center along this vector. Lower values indicate activity closer to (or beyond) the ring center. We performed this procedure for each direction-reward condition. To combine values across direction conditions while maintaining the relationship between reward condition values, we pooled values across rewards within a given reach direction, then z-scored, such that the target preparation axis values for each reach direction overall had zero mean and unit standard deviation (**Fig. 2C**). Results from **Fig. 2C** do not meaningfully change when we use other algorithms to identify the target axes before calculating the target preparation axis (**Fig. S4B**).

We hypothesized that neural activity with smaller projections onto the target preparation axes indicated more poorly-prepared reaches, and hence, would more likely be failures (**Fig. 3A**). To test this, we calculated the average target preparation axis projection (after combining across directions as in **Fig. 2C**) for each reward separately depending on whether the trial was a success, an undershoot, or an overshoot (**Fig. 3D**). To combine across rewards and compare the target preparation axis for successes versus each failure mode, we z-scored values within each reward based on the mean and standard deviation from their successful trials, then pooled across them (**Fig. 3E**). We tested this same procedure using other algorithms to identify the target axes (**Fig. S8**).

*Trial-to-trial variability analysis*

Along with changes in average neural activity as a function of direction and reward, we wanted to assess how trial-to-trial variability was modulated as a function of reward (**Fig. S9**). We calculated trial-to-trial variability within a given reward-direction condition ($\sigma_{d,r}^2$) for each factor of the latent space individually as:

$$\sigma_{d,r}^2 = \frac{1}{T_{d,r}-1} \sum_{t=1}^{T_{d,r}} (x_t - \mu_{d,r})^2 \qquad (3)$$

where $T_{d,r}$ is the number of trials for this reward (*r*) direction (*d*) condition, $x_t$ is the neural activity for trial *t* of this reward-direction condition (in this case, just one dimension of neural activity), and $\mu_{d,r}$ is the average neural activity for reach direction *d* and reward *r*. We then calculated the average trial-to-trial variability for each reward along each factor as the average across reach directions:

$$\sigma_r^2 = \frac{1}{D}\sum_{d=1}^{D}\sigma_{d,r}^2 \qquad (4)$$

where *D* is the total number of reach directions (8 for Monkeys E and R, 4 for Monkey P). This produces a value of trial-to-trial variability for each individual factor of the latent space and each reward condition. To calculate the total trial-to-trial variability within each reward, we took the sum of the trial-to-trial variability across all factors of the latent space.

We then specifically calculated trial-to-trial variability for each reward along specific dimensions of the neural state space that we previously identified, including the reward axis ($\sigma_{r,Rew.Ax.}^2$), target axes ($\sigma_{r,Targ.Ax.}^2$, summed across the two target axes), and axes orthogonal to both of these ($\sigma_{r,Orth.}^2$). The reward axis was orthogonalized to the target axes for this analysis as previously described for Fig. 2A. This yields an exact decomposition of neural variance, such that:

$$\sigma_r^2 = \sigma_{r,Rew.Ax.}^2 + \sigma_{r,Targ.Ax.}^2 + \sigma_{r,Orth.}^2 \qquad (5)$$

Note that, because each animal had a different number of latent factors used for stitching, the number of axes orthogonal to the target and reward axes differed for each animal (Monkey E: 13, Monkey P: 2, Monkey R: 17). To calculate standard error bars for each variance calculation, we performed 1,000 bootstraps within each reward condition and used the standard deviation of the bootstraps.

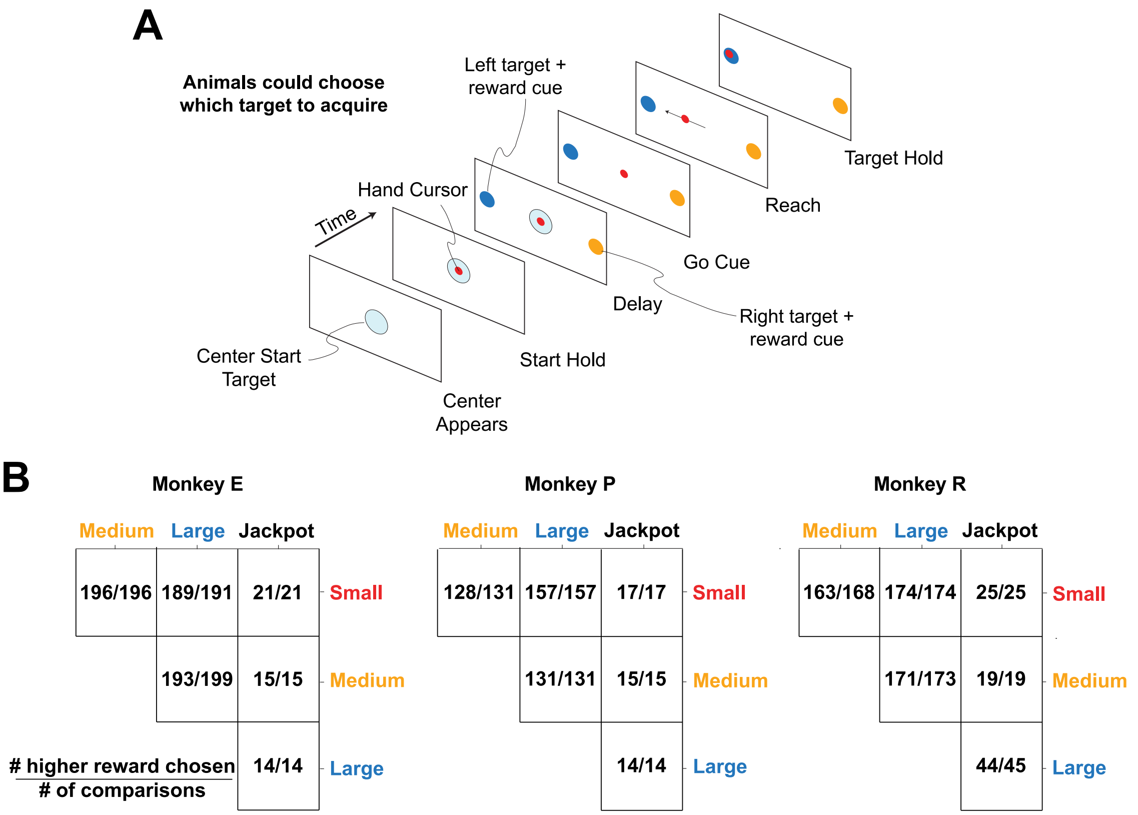


**Fig. S1. A two-target choice task indicates the animals understood the reward values.** All animals performed a choice task identical in structure to the original reaching task but with two diametrically opposed targets presented each trial. This allowed us to assess the animals' understanding of the reward cues by examining their selection of which of the two target to reach to; only one selection was allowed per trial. **(A)** Choice task schematic. Left and right targets are shown for this example, though any pair of diametrically opposed targets from among the positions used in the main task could appear. At the end of a successful trial the animal received the reward corresponding to the cue for the target they acquired. **(B)** Choice behavior for each animal. Results for each possible comparison of rewards is shown. For example, the 1st row shows trials where a Small reward cue was presented at one of the targets, and each column corresponds to the reward cue value of the other target. Each fraction shows the number of trials where the animal selected the higher reward cue target in the numerator with the total number of comparisons for that reward pairing in the denominator. Before data collection, Monkey R initially struggled with selections between Jackpot and other rewards (see Methods, *Behavioral training and recording sessions*). To further assess his understanding of cues, additional Large versus Jackpot reward trials were presented. Given the near-perfect selection of the higher-valued cues, we conclude the animals understood the cues' relative reward values.

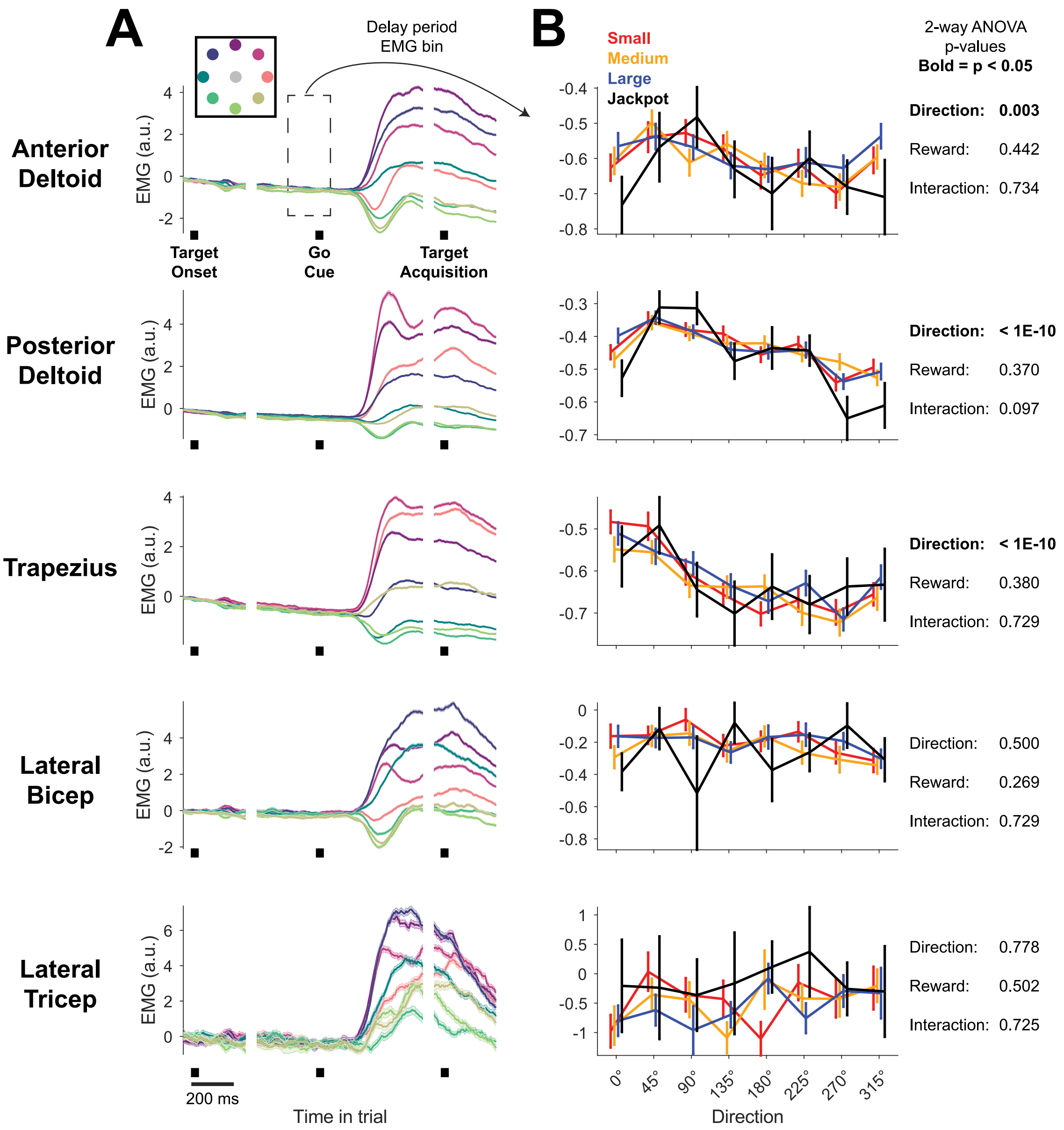


**Fig. S2. Muscle activation during cue presentation does not explain reward-related modulations of neural activity along the reward axis.** To assess if the monotonic "reward axis" effect in neural activity (see **Fig. 1E**) could be explained by increased muscular activation with higher cued rewards before movement onset, we recorded shoulder (anterior deltoid, posterior deltoid, trapezius) and arm (lateral bicep, lateral tricep) surface electromyography (EMG) for Monkey R's working arm during the task (see Methods for EMG processing details). **(A)** EMG signals show strong muscular activation during the trial and directional specificity, as expected. We visualized activation of each muscle as a function of reach direction (see inset for color legend) averaged across successful trials. Shading indicates standard error. Data are aligned to target onset, go cue, and target acquisition, represented by black squares. **(B)** Delay epoch

EMG does not show consistent changes with reward. We calculated the delay period EMG activation as the average of the EMG signal from [-150 50] ms around the go cue minus the baseline for the trial ([-200 0] ms preceding target onset). We then calculated average values and standard errors as a function of direction and reward and plotted them. We used a two-way ANOVA to assess the significance of directional and reward tuning in the delay period EMG signals. While shoulder muscles showed significant directional tuning during the delay period, no muscles showed significant reward tuning, and overall EMG activation in the delay period unsurprisingly far weaker than that during reaching. This provides strong evidence against the view that stiffening of the muscles in the delay period explains the neural effects we have observed along the reward axis. This confirms a previous finding that pre-movement motor cortical representations of reward are unlikely to represent small changes in muscle activity (*26*).

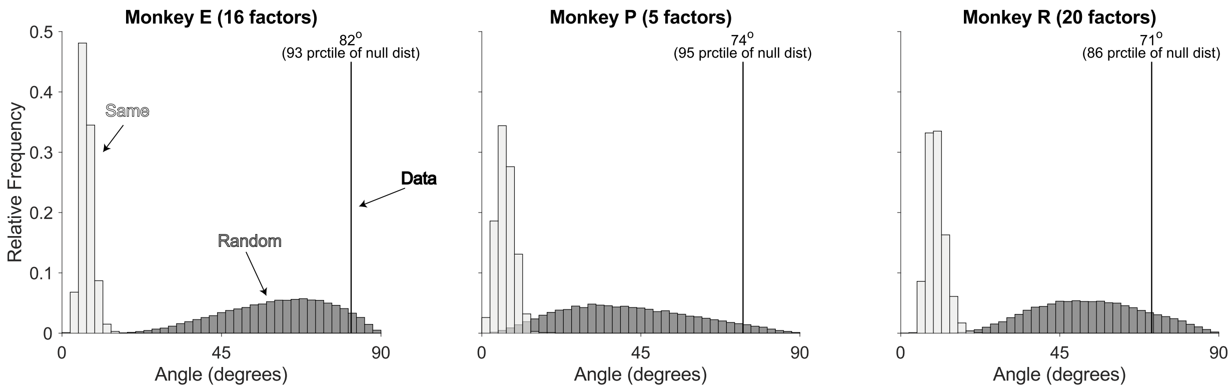


**Fig. S3. The reward axis is nearly orthogonal to the target axes' plane.** We calculated the angle between the reward axis and the 2D plane spanned by the target axes for each animal (black line). Both were found using PCA (see Methods). We also calculated two null distributions for comparison. First, we drew random vectors in the full space of the factor analysis model and measured their angle to the target plane. We did this 50,000 times to construct a distribution of angles between random vectors and the target plane (dark gray, labeled "Random"). Second, we split the data into two halves 1,000 times, randomly apportioning equal numbers of trials for each reward and direction condition. We then found the reward axis in both of these splits and calculated the angle between these two axes (light gray, labelled "Same"). This distribution reflects what the angle between two aligned vectors might be at the resolution of our data. The angles found between the reward axis and target axes' plane were on the upper end of the random vector distribution, far above the aligned distribution, and close to orthogonal.

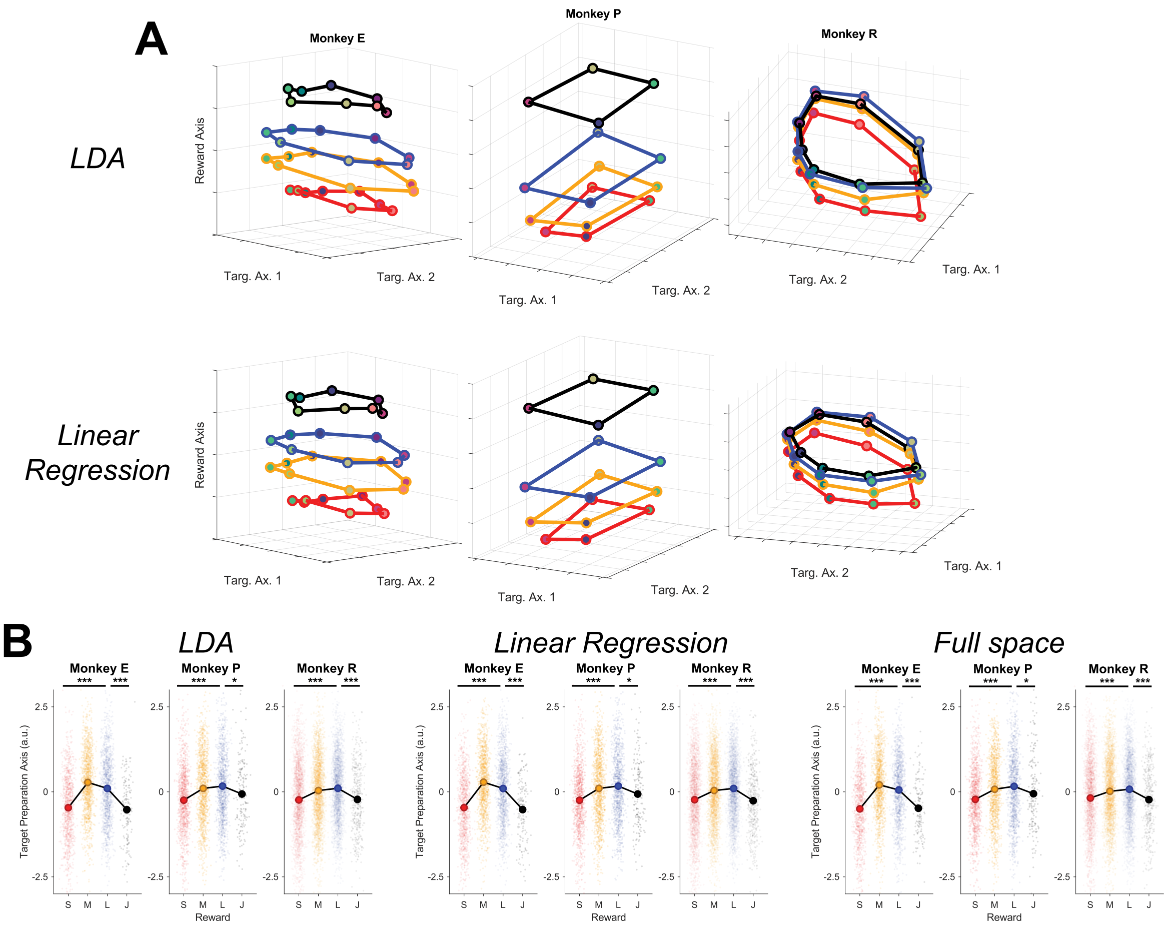


**Fig. S4. Other algorithms reproduce the inverted-U interaction between reward and target information.** Figure formats here correspond to those in Figure 2. **(A)** Along with using PCA to identify dimensions related to target direction information (target axes), we also tested two other algorithms: linear discriminant analysis (LDA) using target location as a categorical label, and linear regression to the target location. For both, we used QR decomposition to orthogonalize the target axes to one another before proceeding. Projections using either of these algorithms replicate the collapse of target conditions for Jackpot rewards. We note that using LDA identifies a plane spanned by the target axes that is closer to orthogonal to the reward axis than the one found using PCA (see Fig. S3 for PCA; for LDA, E: 89.2°, 99.9th percentile of the null distribution, P: 83.5°, 99.0th percentile, R: 86.7°, 99.4th percentile); target axes identified using linear regression were very similar to LDA. **(B)** Quantifying the target preparation axis the same way as described in Figure 2B reproduced the inverted-U revealed as a function of reward for both of these algorithms (LDA and linear regression) for identifying target axes. Further, the target preparation axis calculation and projection does not necessitate projection down to two target axes first and can be effectively calculated from the tuning curves used in the full factor space. We calculated the target preparation axis projections in this way as well and obtained similar results ("Full space").

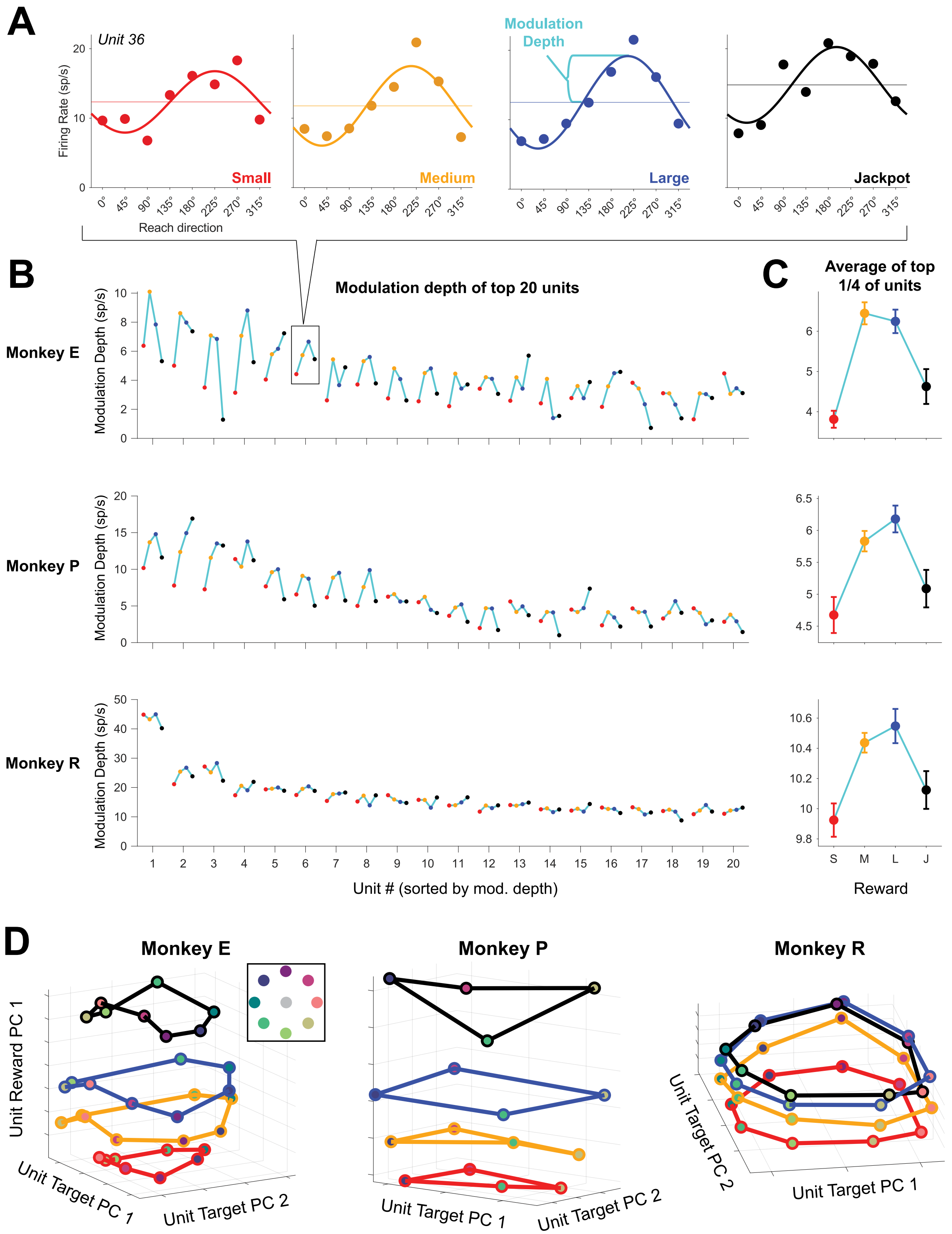


**Fig. S5. Single unit tuning shows an inverted-U relationship between separability of directions with reward.** We wanted to assess the interaction between reward cue and direction encoding in single neural units to compare to the results found at the population level. To do this, we fit cosine functions to the directional tuning of the trial-averaged firing rates of each reach direction (*57*). The phase and amplitude of the cosine model were the parameters fit to the data. We did this within each reward condition for each sorted unit that was present across enough sessions to have at least 10 Jackpot trials for each reach direction (see Methods for how units were identified across sessions). We then calculated each tuning curve's modulation depth, defined as the amplitude of the tuning curve. This provides us with the strength of directional tuning for the given unit under each reward condition. **(A)** Example single unit tuning curves as a function of direction, split by reward condition. The average firing rate as a function of reach direction is shown by the individual points along with the cosine tuning curve fit to those points. **(B)** Modulation depth of the directional tuning curve for each reward for the top 20 units with the greatest Medium-reward modulation depth (connected points come from the same unit). **(C)** Average modulation depth of the ¼ of neurons most tuned to direction reveals that the units with the strongest directional tuning exhibit an inverted-U in tuning strength (depth of modulation) as a function of reward. Error bars are S.E. across units. This reduction in tuning strength with Jackpot rewards shows that the collapse in neural information is evident in the activity of individual neurons. **(D)** We applied PCA on single-unit directional tuning for all available units. Compare to **Fig. 2A** where the collapse in neural information was shown in the neural population space aligned across sessions (see Methods). The similarity between this and Figure 2A indicates that the population level results in the main text are reflected in the responses of individual neurons. From this analysis we can also conclude that the results shown in the main body figures do not result from our stitching algorithm used to combine neural population activity across days.

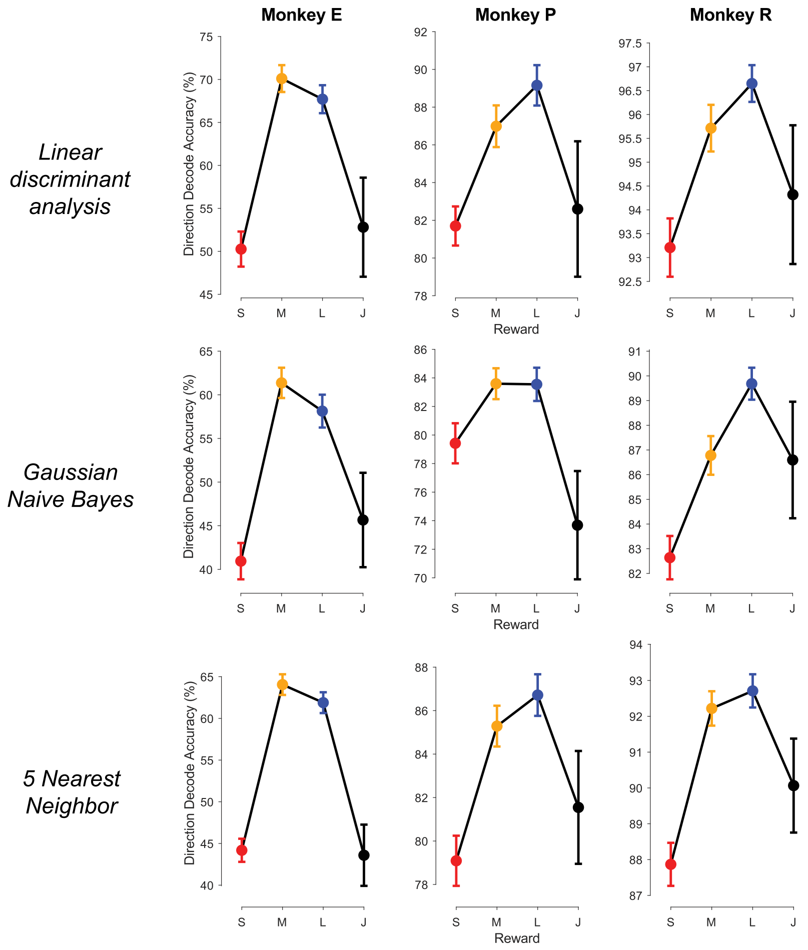


**Fig. S6. Offline decoding of reach direction from neural data exhibits an inverted-U as a function of reward.** Offline decoding is often used to quantify how much information (for example, about target direction) is in a neural signal. Here we use it to show a collapse in neural information induced by Jackpot rewards. To calculate average direction decoding accuracy as a function of reward, we first subsampled (without replacement) all trials for each direction-reward condition down to match the condition with the least number of trials. Then, within each reward, we used 5-fold cross validation to decode cued reach direction from the neural data, training three different types of decoders: linear discriminant analysis (top row), Gaussian Naive Bayes (middle), and 5-nearest neighbors decoder (bottom). We repeated this subsampling, model fitting, and model evaluation procedure 1,000 times and took the average test fold decoding accuracy across the results. We calculated standard error bars within each reward condition using a nested cross-validation method (*58*).

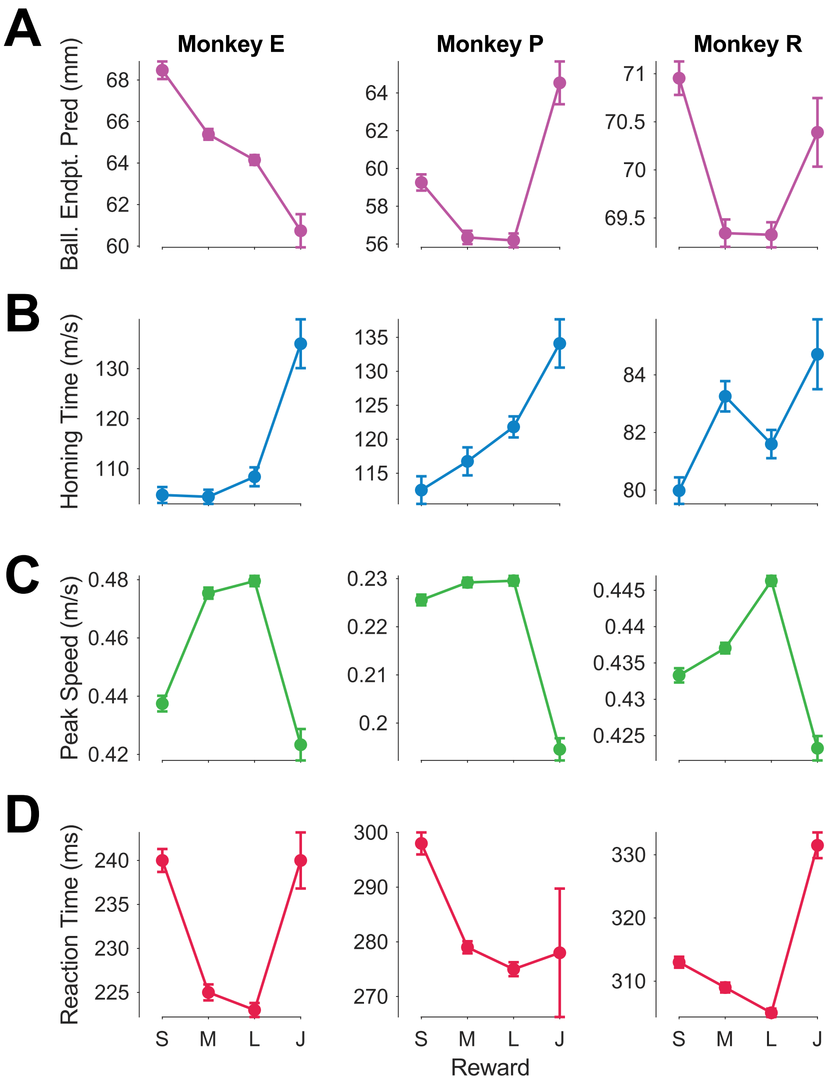


**Fig. S7. Animals exhibit idiosyncratic sources of undershoot failures.** All animals choke under pressure at least in part due to an increased incidence of undershoot failures on Jackpot reward trials (*11*). Animals also undershot more often for Small rewards than for Large rewards. Undershoots can arise through a mixture of three sources: planning a hypometric reach, a slow reaction time, or a slow movement speed. We identified metrics related to each of these causes and evaluated how they depend on reward size. **(A)** Mean ballistic endpoint predictions (± S.E.), predicting where the animal's hand would have landed based on the initial launch of the reach (see Methods). We would predict that planned hypometric reaches would exhibit smaller ballistic endpoint predictions (indicating the ballistic portion of the reach lands closer to the center). Monkey E showed shorter ballistic endpoint predictions with reward, whereas Monkey P showd a U-shaped trend. Monkey R's changes are similar to those of Monkey P, albeit to a much weaker extent. **(B)** Median homing times, defined as the duration it takes the animal to cover the last ⅓ of the distance to the reach target (see Methods). A longer homing time may point to a less ballistic and slower reach, perhaps indicating increased caution in approaching the target. Monkeys E and P show an increase in homing time with reward, especially for Jackpots. Monkey R shows a similar trend, albeit more weakly. **(C)** Mean peak speeds. All animals exhibit an inverted-U in peak speed as a function of reward. However, Monkey P's increase from Small to Large rewards is minute (~2% change). **(D)** Median reaction times (± S.E.). All animals showed faster reaction times with increased rewards up to Large rewards. Monkeys E and R had slower reaction times for Jackpot trials than Large, while Monkey P's were comparable for the two rewards. From these metrics, we conclude that Monkey E's undershoot failures seem to be due to a mixture of slow reaction, slow reaching, and for Jackpots, hypometric reach planning. Monkey P's undershoots for Small rewards seem to be driven by slow reaction times, whereas for Jackpots they are driven by slower reaching. Monkey R shows slower reaction and reaching for both Small and Jackpot rewards. Hence, while all animals undershot the target more often for Small and Jackpot trials than they did for Large rewards, they did so in both subject-specific and reward-specific manners.

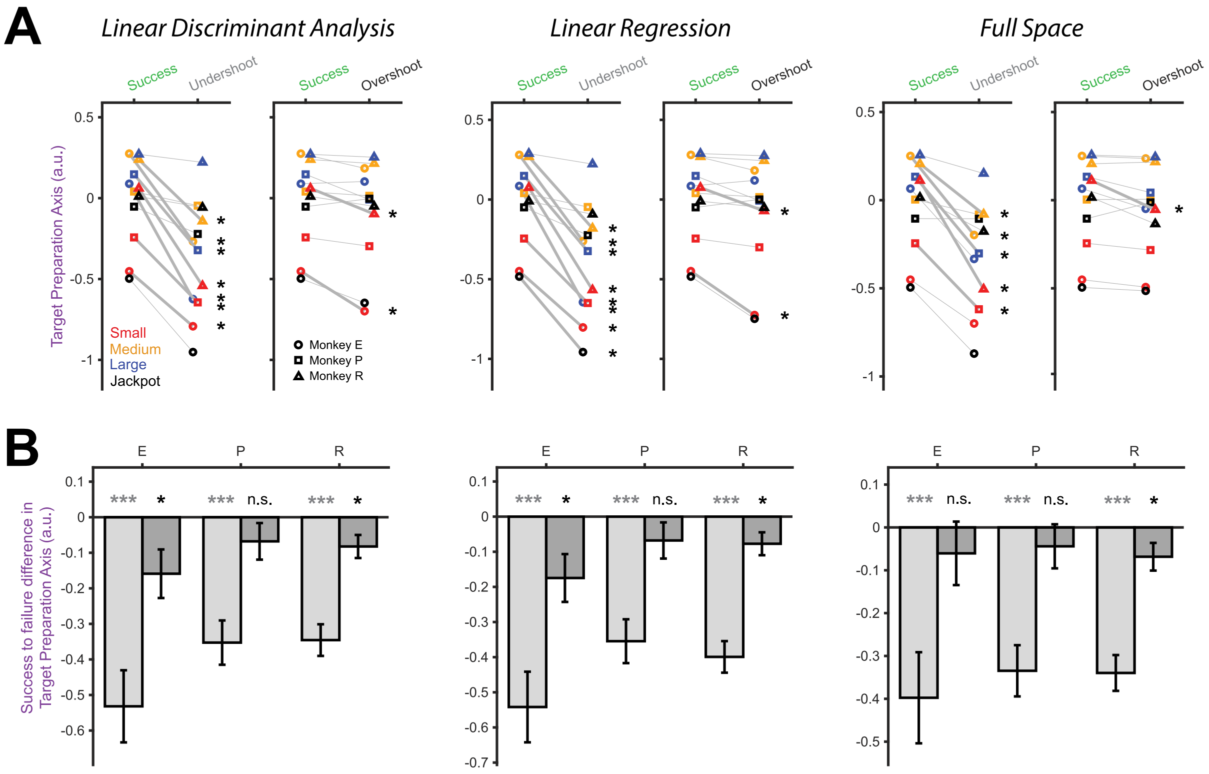


**Fig. S8. The finding of decreased target preparation axis (TPA) projections for undershoot failures does not depend on the algorithm used to find the TPA.** Figure formats here correspond to those in Figure 3. We identified the target preparation axis using four different algorithms: 1. PCA to find target direction encoding axes ("target axes"), 2. linear discriminant analysis (LDA) with direction as a categorical label to identify the target axes, 3. linear regression of neural activity to the target location to define the target axes, 4. not performing any mapping to target axes, but rather simply calculating the target preparation axis on the full space of the data combined across sessions. **(A)** Undershoots consistently showed lower target preparation axis projections than successful trials, regardless of whether we used PCA (see main text Figure 3), LDA (left), linear regression (center), or the full space (right) methods to identify the target preparation axis. Bolded lines and starts indicate statistically significant differences (*$p < 0.05$, Welch's t-test). **(B)** Similarly, we found that these trends hold when combining across rewards (**$p < 0.01$, ***$p < 0.001$, Welch's t-test). We note that for non-PCA algorithms like those shown here, Monkey R's overshoots also show significantly lower target preparation axis value projections ($p < 0.05$, Welch's t-test), and for Monkey E as well if LDA or linear regression are used in the identification of the target axes.

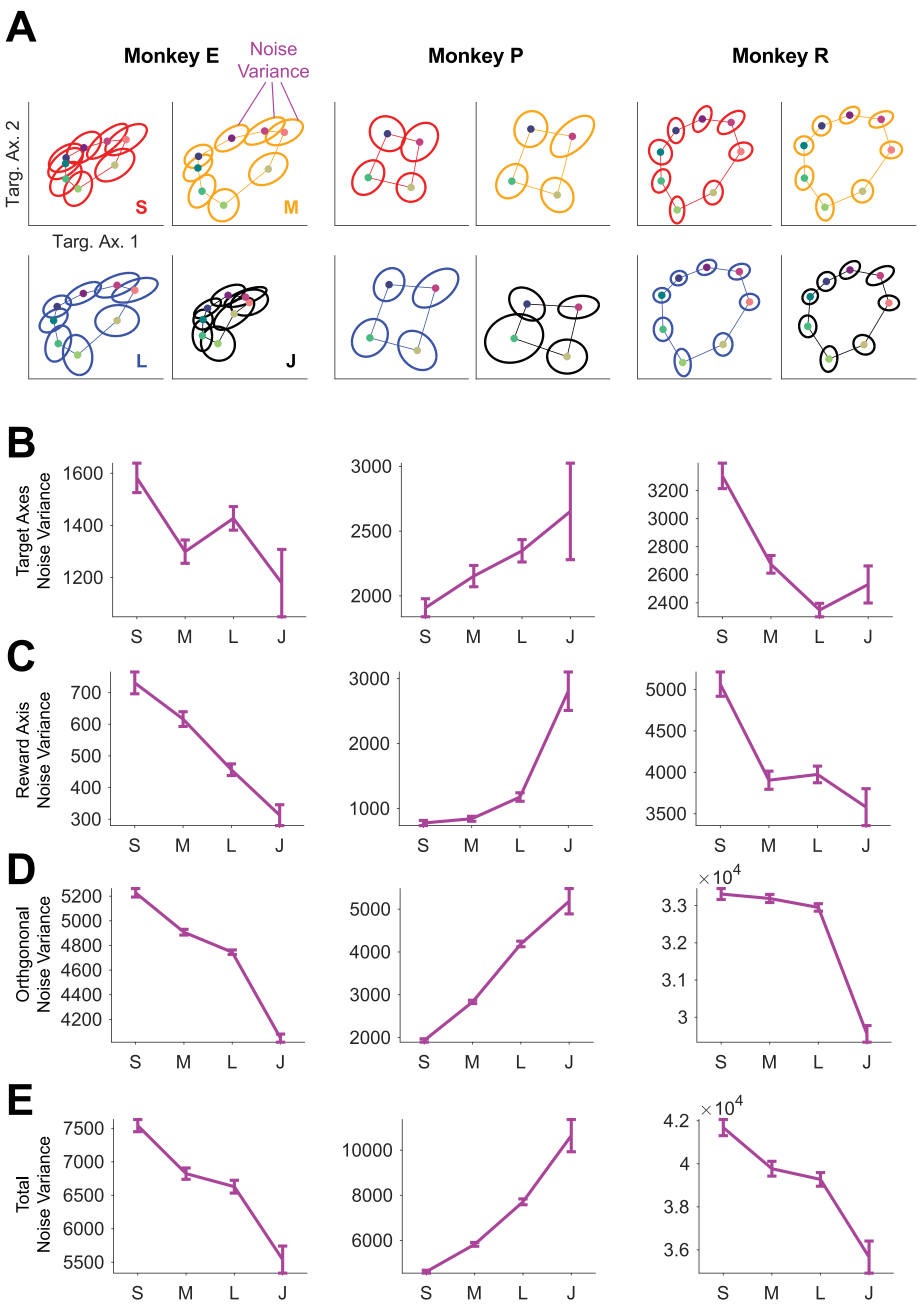


**Fig. S9. Trial-to-trial variability shows inconsistent trends with reward across animals.** To determine if neural noise might also explain choking under pressure, we asked if there were changes in trial-to-trial variability (or "noise variance") about each direction condition average as a function of reward. We calculated the variability about the direction condition means within each reward using a variety of projections and calculated standard error bars using a bootstrapping procedure (see Methods). **(A)** Visualization example of noise variance, shown here in the target axes' plane. Points represent the average neural activity for each reach direction as seen before in Figure 2A. Ellipses indicate the covariance of the single trials' data about the averages. **(B)** Noise variance from the target axes' projection. **(C)** Reward axis noise variance. **(D)** Noise variance in all dimensions orthogonal to the reward and target axes. **(E)** Total noise variance. (Note that we orthogonalized the reward axis with respect to the target plane for this analysis, so that the top row (total) is the sum of the other rows; see Methods). Generally, Monkeys E and R show decreases in variability for greater reward cues, whereas Monkey P shows the opposite. The inconsistent trends in noise variance with reward suggest that reward modulations of noise variability may contribute to choking under pressure but are unlikely to explain the effect in general.

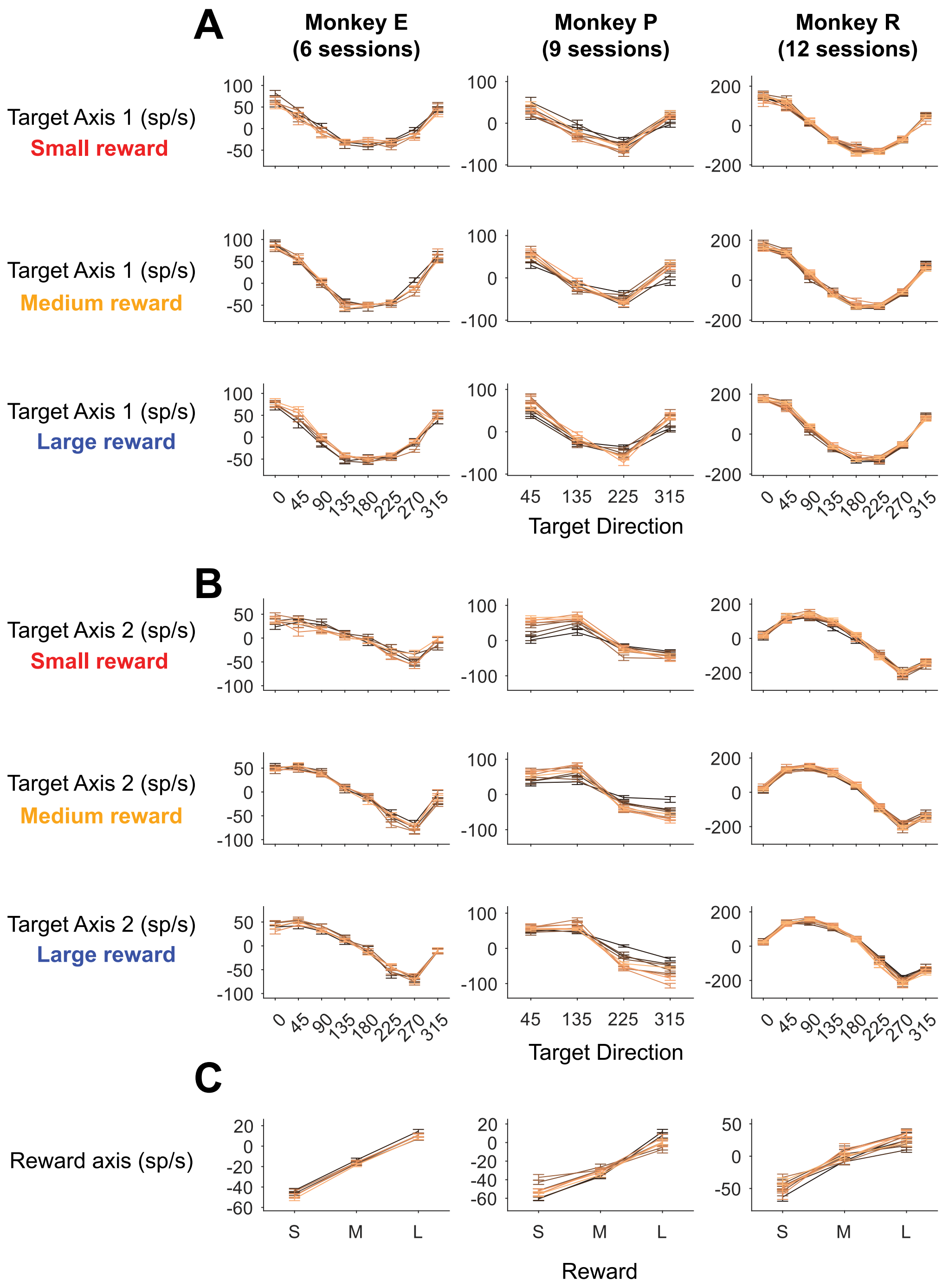


**Fig. S10. Combining neural data across sessions via factor analysis produces stable directional and reward encoding across days.** We combined neural activity across sessions using a modified version of the stabilization algorithm in (*29*) (see Methods). Importantly, the algorithm is unsupervised with respect to direction and reward labels, meaning there is no part of the objective that would require tuning curves across days to be similar. To qualitatively validate that the method worked for our data, we assessed if the neural representations of target direction and reward in the latent state were consistent across days (*55*). Because of low Jackpot trial counts on individual days, we only use Small/Medium/Large reward trials for this analysis. **(A)**

Average projections along the first target axis (left-right target separating). We show individual tuning curves to direction for each day (mean ± S.E.) and for each reward size. Data are presented for each session with earlier sessions being darker colored. **(B)** Similar plots for the second target axis (up-down separating). **(C)** Similar plots for the reward axis.

| **Subject Name** | Monkey E | Monkey P | Monkey R |
|---|---|---|---|
| Array locations | M1 (64 electrodes), PMd (32 electrodes) | M1 (32 electrodes), PMd (64 electrodes) | M1 (192 electrodes) |
| Number of sessions | 6 | 9 | 12 |
| Number of reach target locations | 8 | 4 | 8 |
| Reach target locations | 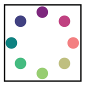 | 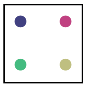 | 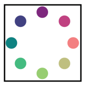 |
| Reach target distance from center (mm) | 85 | 65 | 80 |
| Reach target diameter (mm) | 14.6 | 12 | 12 |
| Center target diameter (mm) | 16.6 | 16 | 18 |
| Cursor diameter (mm) | 6 | 6 | 4 |
| Center hold before target onset (ms) | 200 | [500 600] | 400 |
| Delay period lengths (ms) | [450, 550, 650, 750, 850, 950] | [250 450 650 850] | Drawn uniform random on [200 800] |
| Reach period maximum time | 750 ms | 825 ms | 667 ms |
| Target hold time requirement | 400 ms | 400 ms | 400 ms |
| Small reward size | 0.0 mL | 0.075 mL | 0.0 mL |
| Small reward frequency | 31.67% | 31.67% | 31.67% |
| Small reward cue | 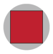 | 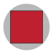 | 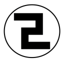 |
| Medium reward size | 0.2 mL | 0.3 mL | 0.22 mL |
| Medium reward frequency | 31.67% | 31.67% | 31.67% |
| Medium reward cue | 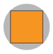 | 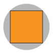 | 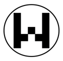 |
| Large reward size | 0.4 mL | 0.525 mL | 0.44 mL |
| Large reward frequency | 31.67% | 31.67% | 31.67% |
| Large reward cue | 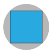 | 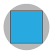 | 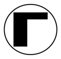 |
| Jackpot reward size | 2.0 mL | 2.4 mL* | 2.2 mL |
| Jackpot reward frequency | 5% | 5% | 5% |
| Jackpot reward cue | 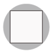 | 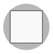 | 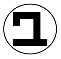 |

**Table S1. Task conditions and experimental details for each animal.** *For session 1 of Monkey P's recording sessions, Jackpot rewards were 4.5 mL due to experimenter error. This was corrected to 2.4 mL for the remaining 8 sessions.

| Reward Tuning Name / Shape | | Reward tuning: Monotonic increase | | | Monotonic decrease | | | U/Inverted-U tuning | | None | Total |
|---|---|---|---|---|---|---|---|---|---|---|---|
| | | S-L-J up | S-L up | J up | S-L-J down | S-L down | J down | U-shape | Inv-U | | |
| 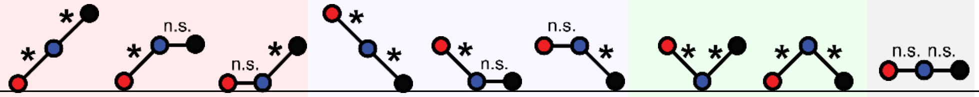 | | | | | | | | | | | |
| **Monkey** | | | | | | | | | | | |
| E | # | 7 | 5 | 4 | 12 | 3 | 5 | 1 | 1 | 4 | **42** |
| | % | 16.7 | 11.9 | 9.5 | 28.6 | 7.1 | 11.9 | 2.4 | 2.4 | 9.5 | |
| | | Inc. Subtotal | | | Dec. Subtotal | | | Subtotal | | | |
| | | | (#) | 16 | | (#) | 20 | (#) | 2 | | |
| | | | (%) | 38.1 | | (%) | 47.6 | (%) | 4.8 | | |
| P | # | 24 | 8 | 22 | 10 | 3 | 6 | 3 | 4 | 33 | **113** |
| | % | 21.2 | 7.1 | 19.5 | 8.8 | 2.7 | 5.3 | 2.7 | 3.5 | 29.2 | |
| | | Inc. Subtotal | | | Dec. Subtotal | | | Subtotal | | | |
| | | | (#) | 54 | | (#) | 19 | (#) | 7 | | |
| | | | (%) | 47.8 | | (%) | 16.8 | (%) | 6.2 | | |
| R | # | 49 | 53 | 7 | 19 | 23 | 14 | 4 | 13 | 122 | **304** |
| | % | 16.1 | 17.4 | 2.3 | 6.3 | 7.6 | 4.6 | 1.3 | 4.3 | 40.1 | |
| | | Inc. Subtotal | | | Dec. Subtotal | | | Subtotal | | | |
| | | | (#) | 109 | | (#) | 56 | (#) | 17 | | |
| | | | (%) | 35.9 | | (%) | 18.4 | (%) | 5.6 | | |
| **Total** | # | 80 | 66 | 33 | 41 | 29 | 25 | 8 | 18 | 159 | **459** |
| | % | 17.4 | 14.4 | 7.2 | 8.9 | 6.3 | 5.4 | 1.7 | 3.9 | 34.6 | |
| | | Inc. Subtotal | | | Dec. Subtotal | | | Subtotal | | | |
| | | | (#) | 179 | | (#) | 95 | (#) | 26 | | |
| | | | (%) | 39.0 | | (%) | 20.7 | (%) | 5.7 | | |

**Table S2. Single unit reward tuning statistics.** S-L-J indicates trends present from Small to Large and Large to Jackpot. J up / down indicates Jackpots versus smaller rewards were the only significant difference. "Inc. Subtotal" indicates the sum of neurons with "Monotonic increase" tuning. Similar subtotals are shown for the "Monotonic decrease" and "U/Inverted-U tuning" sections.

**Movie S1.** Rotations of Figure 2A to show the 2D projection spanned by the reward and target axes. Small, Medium, and Large reward are shown in the top row, while Large and Jackpot are shown in the bottom row.

### Supplemental material references


11. A. L. Smoulder, N. P. Pavlovsky, P. J. Marino, A. D. Degenhart, N. T. McClain, A. P. Batista, S. M. Chase, Monkeys exhibit a paradoxical decrease in performance in high-stakes scenarios. *Proc. Natl. Acad. Sci. U.S.A.* **118**, e2109643118 (2021).

48. M. D. Golub, B. M. Yu, S. M. Chase, Internal models for interpreting neural population activity during sensorimotor control. *eLife*. **4**, e10015 (2015).

49. W. E. Bishop, B. M. Yu, "Deterministic symmetric positive semidefinite matrix completion" in *Advances in Neural Information Processing Systems* (2014).

50. C. Pandarinath, D. J. O'Shea, J. Collins, R. Jozefowicz, S. D. Stavisky, J. C. Kao, E. M. Trautmann, M. T. Kaufman, S. I. Ryu, L. R. Hochberg, J. M. Henderson, K. V. Shenoy, L. F. Abbott, D. Sussillo, Inferring single-trial neural population dynamics using sequential auto-encoders. *Nat Methods*. **15**, 805–815 (2018).

51. J. Jude, M. G. Perich, L. E. Miller, M. H. Hennig, Robust alignment of cross-session recordings of neural population activity by behaviour via unsupervised domain adaptation (2022), (available at http://arxiv.org/abs/2202.06159).

52. W. E. Bishop, thesis, Carnegie Mellon University (2015).

53. M. Nonnenmacher, S. C. Turaga, J. H. Macke, "Extracting low-dimensional dynamics from multiple large-scale neural population recordings by learning to predict correlations" in *Advances in Neural Information Processing Systems* (2017).

54. G. W. Fraser, A. B. Schwartz, Recording from the same neurons chronically in motor cortex. *Journal of Neurophysiology*. **107**, 1970–1978 (2012).

55. J. A. Gallego, M. G. Perich, R. H. Chowdhury, S. A. Solla, L. E. Miller, Long-term stability of cortical population dynamics underlying consistent behavior. *Nat Neurosci*. **23**, 260–270 (2020).

56. B. M. Yu, J. P. Cunningham, G. Santhanam, S. I. Ryu, K. V. Shenoy, M. Sahani, Gaussian-process factor analysis for low-dimensional single-trial analysis of neural population activity. *Journal of Neurophysiology*. **102**, 614–635 (2009).

57. A. Georgopoulos, J. Kalaska, R. Caminiti, J. Massey, On the relations between the direction of two-dimensional arm movements and cell discharge in primate motor cortex. *J. Neurosci.* **2**, 1527–1537 (1982).

58. S. Bates, T. Hastie, R. Tibshirani, Cross-validation: what does it estimate and how well does it do it? (2022), (available at http://arxiv.org/abs/2104.00673).
